# A Novel MHC-Independent Mechanism of Tumor Cell Killing by CD8^+^ T Cells

**DOI:** 10.1101/2023.02.02.526713

**Authors:** Emily Lerner, Karolina Woroniecka, Vincent D’Anniballe, Daniel Wilkinson, Selena Lorrey, Jessica Waibl-Polania, Lucas Wachsmuth, Alexandra Miggelbrink, Jude Raj, Aditya Mohan, Sarah Cook, William Tomaszewski, Xiuyu Cui, Mustafa Khasraw, Michael D. Gunn, Peter E. Fecci

## Abstract

The accepted paradigm for both cellular and antitumor immunity relies upon tumor cell kill by CD8^+^ T cells recognizing cognate antigens presented in the context of target cell major histocompatibility complex class I (MHC I) molecules. Likewise, a classically described mechanism of tumor immune escape is tumor MHC-I downregulation. Here, we report that CD8^+^T cells maintain the capacity to kill tumor cells that are entirely devoid of MHC-I expression. This capacity proves to be dependent on interactions between T cell NKG2D and tumor NKG2D ligands (NKG2DL). Necessarily, tumor cell kill in these instances is antigen-independent, although prior T cell antigen-specific activation is required and can be furnished by myeloid cells or even neighboring MHC-replete tumors cells. These mechanisms are active *in vivo* in mice, as well as *in vitro* in human tumor systems, and are obviated by NKG2D knockout or blockade. Tumor cell killing following T cell NKG2D engagement is Fas-independent and appears to involve granzyme. These studies potentially obviate the long-advanced notion that downregulation of MHC-I is a viable means of tumor immune escape, and instead identify the NKG2D/NKG2DL axis as a novel therapeutic target for enhancing T cell-dependent anti-tumor immunity against MHC loss variants.

## Introduction

The long-accepted paradigm for adaptive anti-tumor cellular immunity relies on antigen-specific tumor targeting by activated CD8^+^ T cells. CD8^+^ T cell cytotoxicity, in turn, is classically believed to depend upon T cell receptor (TCR) recognition of tumor antigens presented exclusively in the context of cell surface major histocompatibility complex I (MHC-I) molecules. A critical component to the antigen presentation function of MHC-I is beta-2-microglobulin (β2m), which is conserved across all classical MHC-I alleles present in mice and humans. In turn, mutations in β2m leading to decreased or absent MHC-I expression are purported to constitute a common mechanism by which tumors are able to evade T cell responses, rendering them immunologically “cold” [1–4]. The frequency of such MHC-I downregulation varies by tumor histology, with certain tumors, such as glioblastoma (GBM), often expressing little to no MHC-I [5, 6].

Recent pre-clinical and clinical studies, however, have demonstrated somewhat mixed roles for MHC-I and β2m in dictating responses to cancer immune-based platforms, such as immune checkpoint blockade (ICB). While some have suggested that resistance to ICB emerges through inactivation of tumor antigen presentation [7–9], for instance, others have implied that low tumor β2m and MHC-I expression are instead associated with favorable prognosis [10, 11]. These mixed findings highlight the need for further investigation into the role of tumor MHC-I, as well as revisiting traditional notions of anti-tumor immunity.

Following our recent investigations into T cell exhaustion and ICB-resistance in tumors [12, 13] we sought to better evaluate the role of MHC-I expression within the tumor microenvironment (TME). Herein, we reveal that T cell-activating immunotherapies remain effective against glioma and melanoma lines engineered to lack MHC-I expression. Such efficacy proves independent of natural killer (NK) cells and, instead, relies consistently upon CD8^+^ T cell-mediated cytotoxicity, despite the absence of MHC-I on tumor targets. This cytotoxicity is both antigen- and MHC-agnostic and, instead, is mediated by T cell NKG2D engagement of non-classical MHC-I (NKG2D ligands) on tumor cells. Subsequent tumor cell kill depends on prior TCR activation (albeit even by irrelevant antigen) and appears to involve granzyme and/or perforin, but is not dependent on Fas. This novel mechanism of cytotoxicity is active *in vivo* in mice, as well as *in vitro* in human cells, and is required for killing of MHC-negative tumor cells, even in tumors with heterogenous MHC expression. Such findings challenge the traditional model of T cell-mediated tumor cell kill and likewise provide a novel therapeutic blueprint for licensing immune responses in MHC-loss tumor cell variants.

## Results

### The Efficacy of Immunotherapy Against Tumors Lacking MHC-I Still Depends on CD8^+^ T Cells

We have previously demonstrated the efficacy of combination 4-1BB agonism and anti-PD-1 checkpoint blockade (aPD-1/4-1BB) in a murine CT2A glioma model. Unsurprisingly, such efficacy proved dependent on the presence of CD8^+^ T cells [12]. More recently, to examine the antigen-specific response, we engineered a CT2A mouse glioma tumor line to express tyrosinase-related peptide 2 (TRP-2), a weakly immunogenic model antigen, creating CT2A-TRP2. Combination aPD-1/4-1BB therapy demonstrated similar efficacy against orthotopically implanted CT2A-TRP2, resulting in 80% long term survival (**Supplementary Fig. 1A).**

With initial intent to examine the impact of tumor antigen presentation on facets of the TME, we utilized CRISPR to knockout (KO) β2m in the CT2A-TRP2 line, generating CT2A-TRP2-β2mKO (**Figure 1A**). CT2A-TRP2-β2mKO lacked MHC-I expression (H-2K^b^ and H-2D^b^) by flow cytometry. MHC-I KO was further confirmed with an interferon-gamma (IFN-γ) stimulation assay [13]. IFN-γ stimulation resulted in upregulation of H-2K^b^ and H-2D^b^ in parental CT2A-TRP2, but not in CT2A-TRP2-β2mKO (**Figure 1B**). Additionally, TRP-2 TCR-transduced CD8^+^ T cells (TRP2 T cells) failed to kill CT2A-TRP2-β2mKO tumors *in vitro*, but efficiently killed parental CT2A-TRP2 tumors, providing functional validation of the MHC-I knockdown in the CT2A-TRP2-β2mKO line (**Figure 1C)**.

**Figure 1:**
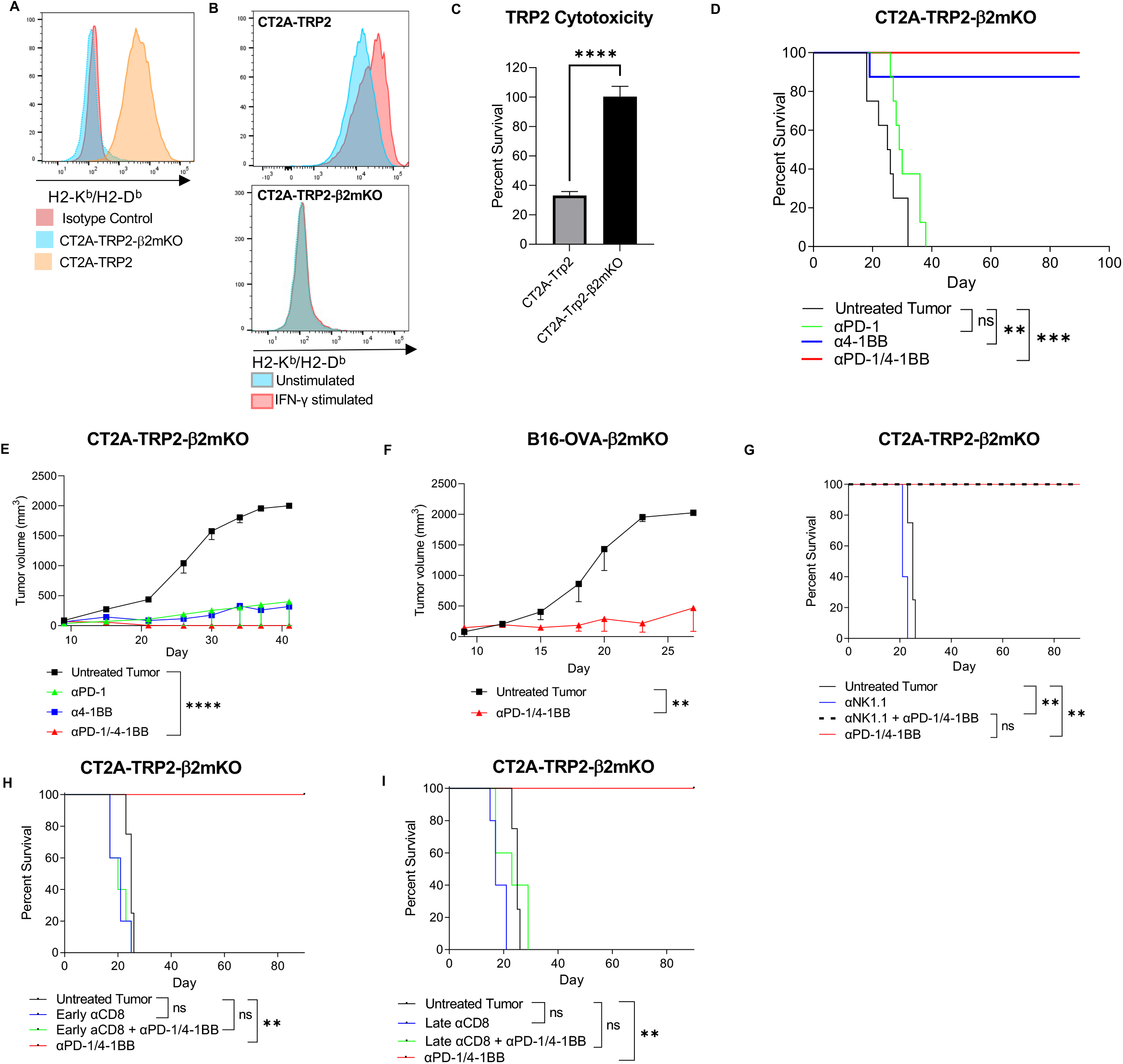
Immunotherapies remain effective against tumor cells lacking MHC-I in a CD8^+^ T cell-dependent manner. **A**. Flow cytometry of tumors cells stained for H2-K^b^ and H2-D^b^, the HLA haplotypes expressed in C57Bl/6 mice **B.** Flow cytometry of tumors stained for H2-K^b^ and H2-D^b^ following 24 hr *in vitro* stimulation with IFN-γ (20 ng/mL). **C.** Percent survival of CT2A-TRP2 tumors or CT2A-TRP2-β2mKO tumors co-cultured with tumor specific TRP-2-specific CD8^+^ T cells (TRP2 T cells), 10:1 ET. After 24 hrs of co-culture, the number of remaining viable tumor cells was quantified using flow cytometry and percent survival was calculated in comparison to tumor only wells. **D**. Kaplan-Meier survival curve of C57BL/6 mice implanted with CT2A-TRP2-β2mKO IC then treated with αPD-1, α4-1BB, αPD-1 +α4-1BB (αPD-1/4-1BB), or PBS vehicle control. N=8/ group. **E-F** Subcutaneous tumor growth curve of CT2A-TRP2-β2mKO (**e**) and B16-OVA-β2mKO (**f**) implanted into C57BL/6. N=5/group. **G**. Kaplan-Meier survival curve of mice implanted with CT2A-TRP2-β2mKO IC in the presence or absence of NK depletion with αNK1.1. N=5/group. **H,I.** Kaplan-Meier survival curve of mice implanted with CT2A-TRP2-β2mKO IC followed by CD8^+^T depletion with αCD8α in the early priming stage (**h**) (day 3 after tumor implantation) or later effector stage (**i**) (day 7 after tumor implantation). Data in **c** are shown as mean ±s.e.m., 6 technical replicates, *P* value was calculated with a two-tailed student’s T test. Tumor growth curves in **e** and **f** were compared with two-way repeated measures ANOVA. Survival in **d,g-i** was assessed by Gehan-Breslow-Wilcoxon test, with Bonferroni correction for multiple comparisons. \**P* < 0.05, \*\**P* < 0.01, \*\*\**P* < 0.001, \*\*\*\**P* < 0.0001. NS: not significant. ET: Effector to target ratio.

Surprisingly, we found that mice harboring intracranial (IC) CT2A-TRP2-β2mKO tumors remained responsive to immunotherapy, demonstrating 100% long-term survival in response to combination aPD-1/4-1BB treatment, and 90% long-term survival in response to 4-1BB monotherapy (**Figure 1D**). As with CT2A-TRP2, the CT2A-TRP2-β2mKO tumor was uniformly fatal without treatment. Immunotherapeutic efficacy against CT2A-TRP2-β2mKO was not specific to the IC compartment, as aPD-1/4-1BB remained effective against this MHC-I negative glioma when implanted subcutaneously (SC) as well (**Figure 1E**). Furthermore, aPD-1/4-1BB retained efficacy against orthotopically implanted melanomas lacking MHC-I (B16-OVA-β2mKO) **(Figure 1F).** Ultimately, immunotherapeutic success in MHC-I-negative tumors was restricted neither to gliomas, nor to the IC compartment.

Intrigued, we sought to determine the immune cell population responsible for the efficacy of immunotherapy against MHC-I-negative tumors. As NK cells are known to eliminate cells with absent or reduced MHC [14], we first investigated by depleting NK cells prior to implanting CT2A-TRP2-β2mKO tumors (NK depletion depicted in **Supplementary Figure 1B)**. Even in the absence of NK cells, aPD-1/4-1BB still elicited 100% long-term survival (**Figure 1G**). We therefore explored whether immunotherapeutic efficacy against CT2A-TRP2-β2mKO might still be dependent on CD8^+^ T cells, even in the absence of tumor MHC-I expression. Depletion of CD8^+^T cells (**Supplementary Figure 1C**) completely abrogated the survival benefit of aPD-1/4-1BB in CT2A-TRP2-β2mKO. This proved true whether CD8^+^ T cells were depleted prior to tumor implantation (**Supplementary Figure 1D**), or at early (Day 3) or late (Day 7) timepoints post-tumor implantation (**Figures 1H-I**). Of note, CD4^+^ T-cell depletion had a modest but nonsignificant impact on survival **(Supplementary Figures 1E-F)**.

### Antigen Specific Killing Persists in the Absence of Tumor MHC-I

Continued dependence on CD8^+^ T cells raised the question as to whether cytotoxic killing of MHC-I-negative tumors still occurs in an antigen-specific fashion. Interestingly, we noted that αPD-1/4-1BB-treated mice bearing MHC-I-negative CT2A-TRP2-β2mKO tumors accumulated a higher number of TRP-2-specific CD8^+^ T cells in the tumor compared to likewise-treated mice bearing MHC-I-positive CT2A-TRP2 **(Figure 2A,** tetramer staining shown in **Supplementary Figure 2A)**. To test then the role of antigen-specificity, we implanted CT2A-TRP2-β2mKO or CT2A-β2mKO (lacking both TRP-2 and MHC-I expression) IC into mice lacking CD8^+^ T cells (CD8KO). Mice subsequently received an adoptive transfer of TRP-2-specific CD8^+^ T cells (TRP-2 ALT), producing a T cell compartment exclusively specific for TRP-2 (schematic in **Figure 2B).** TRP-2 ALT in combination with aPD-1/4-1BB was sufficient to eliminate 100% of CT2A-TRP2-β2mKO tumors, (**Figure 2C**) but was uniformly unsuccessful against CT2A-β2mKO tumors lacking TRP-2 expression (**Figure 2D**). Additionally, we implanted CT2A-TRP2-β2mKO tumors IC into transgenic OT-1 mice. The majority of T cells in these mice recognize the ovalbumin peptide residues 257-264 (OVA257-264; SIINFEKL) in the context of H2K^b^, but possess no reactivity against TRP-2. Combination aPD-1/4-1BB was completely ineffective against CT2A-TRP2-β2mKO in these mice (**Figure 2E**), further suggesting that tumor-specific T cells remain necessary, even in the absence of tumor MHC-I antigen presentation.

**Figure 2:**
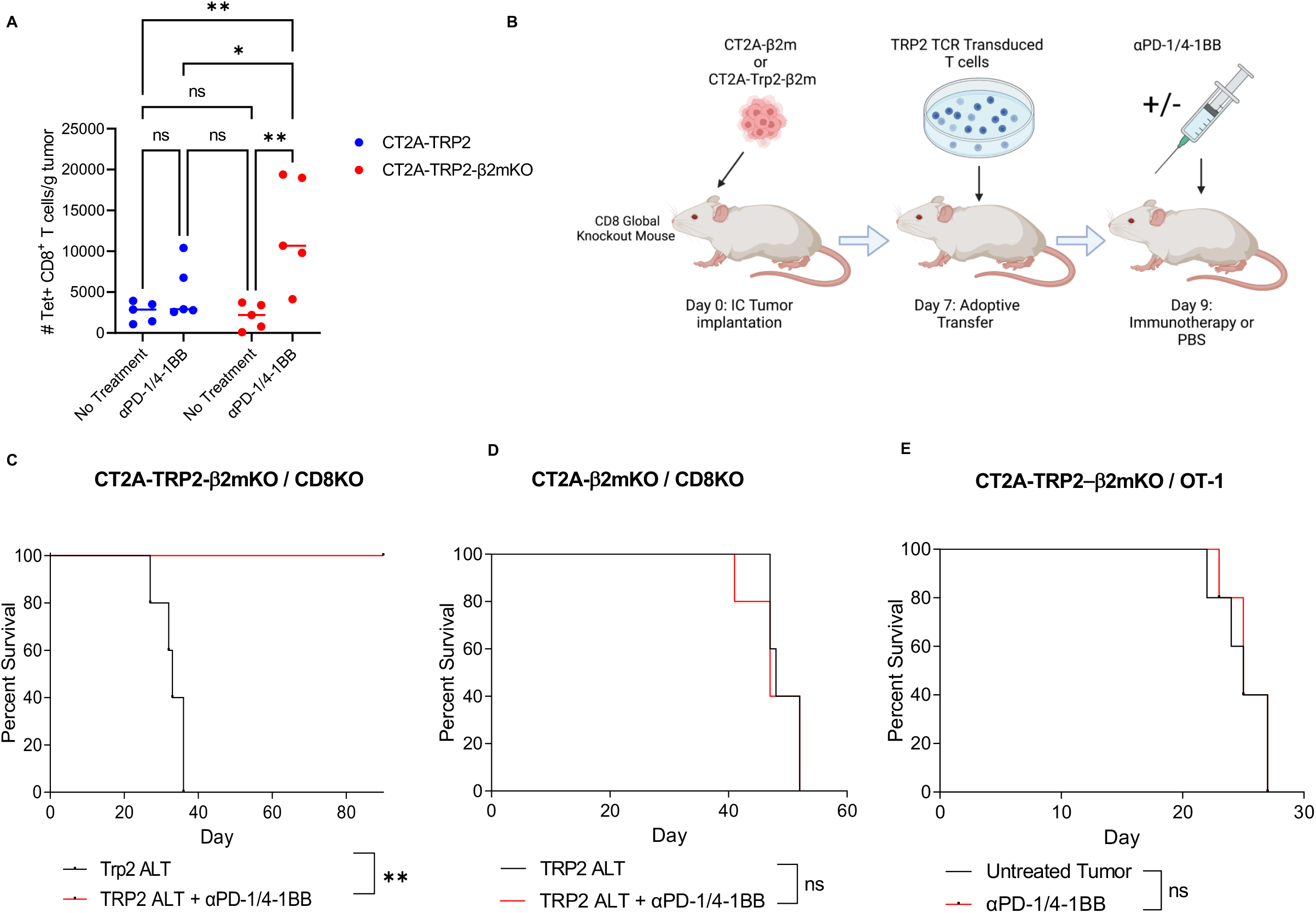
Antigen-specific killing persists in the absence of tumor MHC-I. **A.** Absolute count of tumor-specific TRP2 CD8^+^ T cells from C57BL/6 mice bearing either IC CT2A-TRP2 or CT2A-TRP2-β2mKO tumors. Mice were implanted with CT2A-TRP2-β2mKO or CT2A-TRP2 IC then treated with intraperitoneal αPD-1/4-1BB at day 18. Tumors were then harvested 24 hours later, dissociated, TRP-2 tetramer stained, and assessed by flow cytometry. **B**. Schematic of experimental design. Kaplan-Meier survival curve of CD8^+^ T cell global knockout mice (CD8KO) implanted with IC CT2A-TRP2-β2mKO (**c**) or CT2A-β2mKO (**d**) (lacking TRP-2 expression). Mice were treated with TRP2 TCR engineered T cells on day 7, followed by IP αPD-1/4-1BB therapy or vehicle control. **E**. Kaplan-Meier survival curve of OT-1 mice implanted with CT2A-TRP2-β2mKO tumors, which do not express the OVA antigen, creating a model with little to no tumor specific T cells present. Data in **a** are shown as mean ±s.e.m. *P* value was calculated with one-way ANOVA with post hoc Tukey’s test. N=5/group Survival in **c-e** was assessed by Gehan-Breslow-Wilcoxon test. *P* values were Bonferroni corrected for multiple comparisons. Schematic made with Biorender.com.

### The Efficacy of Immunotherapy Against MHC-I-Negative Tumors Requires Antigen-Specific Interaction Between CD8^+^ T Cells and Antigen Presenting Cells, but Not Between CD8^+^ T Cells and Tumor

Above we demonstrated that TRP-2 T cells failed to kill CT2A-TRP2-β2mKO tumor cells *in vitro*(**Figure 1C**), confirming that CD8^+^ T cells *alone* cannot kill MHC-I-negative tumors, even in the presence of a cognate antigen. We hypothesized then that an additional cell population must be facilitating the observed cytotoxicity *in vivo. In vivo*, infiltrating myeloid cells constitute the bulk of immune cells within the brain tumor TME [15]. We therefore examined whether adding bone marrow-derived macrophages (BMDMs) would license *in vitro* killing of CT2A-TRP2-β2mKO tumor by TRP-2 T cells. CT2A-TRP2 or CT2A-TRP2-β2mKO (**Figure 3A**) tumor cells were cultured with TRP-2 T cells in the presence of either TRP-2 peptide-loaded BMDMs (TRP-2 Mφ) or unpulsed BMDMs (Unpulsed Mφ). While TRP-2 T cells alone (but not TRP-2 Mφ) efficiently killed MHC-I-positive CT2A-TRP2, neither TRP-2 T cells alone nor TRP-2 Mφ alone proved capable of killing MHC-I-negative CT2A-TRP2-β2mKO. Remarkably, however, the combination of TRP-2 Mφ and TRP-2 T cells was indeed sufficient to kill CT2A-TRP2-β2mKO *in vitro*.Interestingly, the combination of unpulsed Mφ and TRP-2 T cells also failed to result in significant tumor cell kill, suggesting that antigen-cognate interactions between Mφ and CD8^+^ T cells are necessary for deriving cytotoxicity against tumor cells lacking MHC-I. Sham TCR-transduced CD8^+^ T cells failed to kill either tumor cell line, even in the presence of TRP-2 loaded Mφ. Importantly, we recapitulated these findings in an additional murine glioma line (GL261) expressing a different target antigen (OVA) using OVA-specific OT-1 T cells and OVA-pulsed MΦ (OVA MΦ) (**Figure 3B-D**).

**Figure 3:**
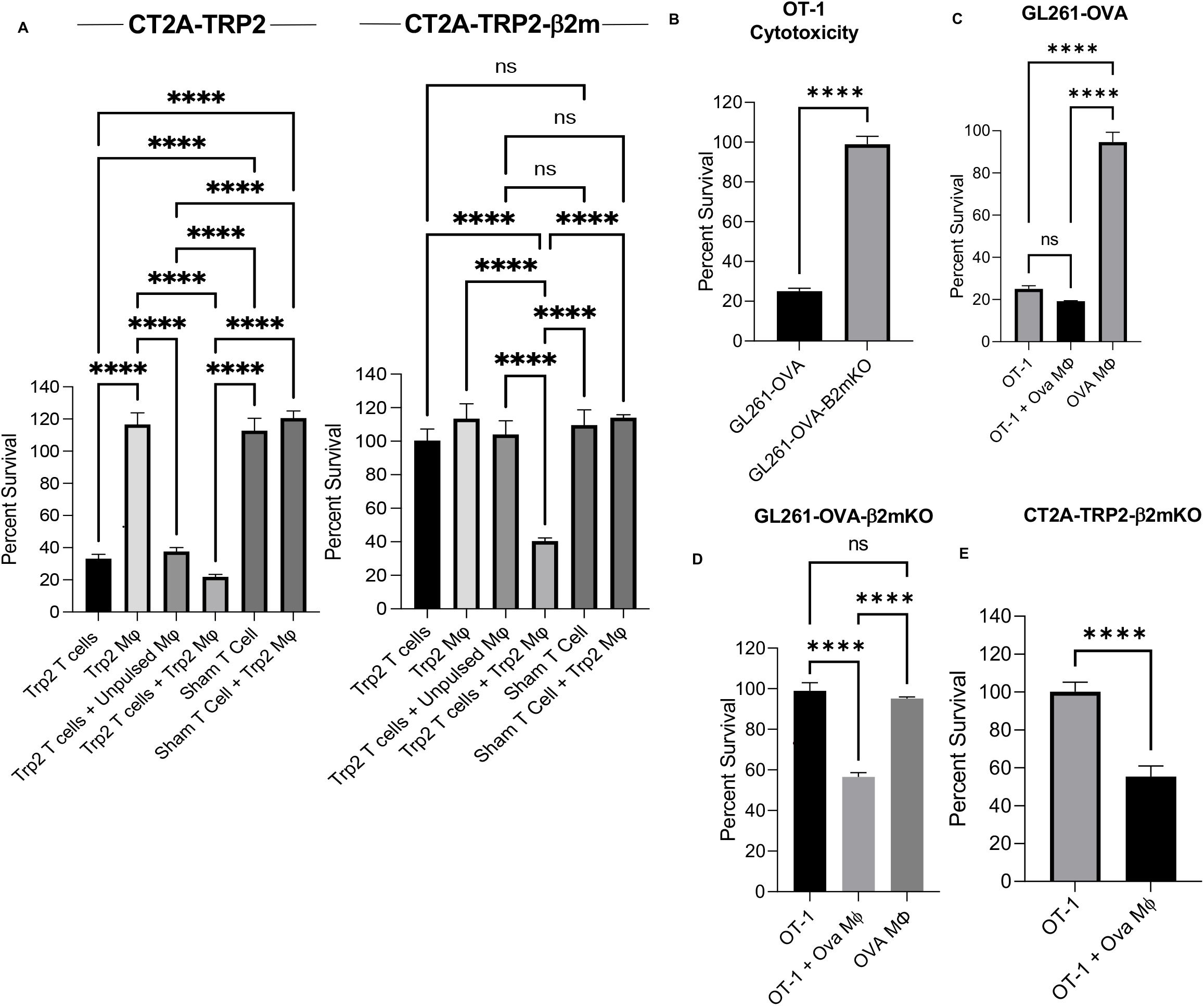
In the absence of tumor MHC-I, CD8^+^ T cell killing requires antigen-specific activation by antigen presenting cells *in vitro*. **A**. Percent survival of CT2A-TRP2 tumors or CT2A-TRP2-β2mKO tumors co-cultured with TRP2 T cells and macrophages. Bone marrow derived macrophages (BMDMs) were loaded with TRP-2 peptide (180-188) (TRP2 Mϕ) or vehicle control (unpulsed Mϕ). TRP2 T cells were generated as previously described and isolated based on CD8 expression. TRP2 T cells or sham transduced T cells (Sham T cells) were co-cultured with TRP2 Mϕ, unpulsed Mϕ and CellTrace™ Violet stained tumor cells at a 10:1 ET. After 24 hr co-culture, the number of remaining viable tumor cells was quantified using flow cytometry and percent survival was calculated in comparison to tumor only wells. **B.** OVA SIINFEKL peptide (257-264) specific OT-1 T cells were co-cultured with a murine glioma model expressing OVA antigen (GL261-OVA) or MHC-I-negative GL261-OVA-β2mKO for 24 hours at a 10:1 ET. Percent survival was calculated as described above. **C,D**. BMDMs were loaded with OVA SIINFEKL peptide (257-264) (OVA Mϕ) and stained with CellTrace™ CFSE. GL261-OVA tumors (**c**) or GL261-OVA-β2mKO tumors (**d**) were then co-cultured with OT-1 T cells and/or OVA Mϕ and percent survival after 24 hours was calculated. **E.** CT2A-TRP2-β2mKO tumors were co-cultured with non-tumor specific OT-1 CD8^+^ T cells (OT-1 T cells) or OT-1 T cells and CFSE stained OVA Mϕ at a 10:1 total ET. All data are shown as mean ±s.e.m. *P* values were determined by two-tailed, unpaired Student’s t-test for **b & e,** and one-way ANOVA with post-hoc Tukey’s test for **a**, **c & d**. Data in **a-f** are representative findings of 6 technical replicates from one of a minimum of two independently repeated experiments with similar results.

While these results suggested the importance of antigen-cognate interactions between antigen presenting cells (APCs) and T cells, the role of such interactions between T cells and tumor cells lacking MHC-I remained less clear. Therefore, we performed *in vitro* cytotoxicity experiments using antigen-matched OT-1 CD8^+^ T-cells and OVA Mφ co-cultured with OVA-negative CT2A-TRP2-β2mKO tumor cells. We found that the combination of OT-1 T cells and OVA Mφ was sufficient to kill CT2A-TRP2-β2mKO tumor cells, even though the tumor itself expressed neither MHC-I, nor the cognate OVA antigen (**Figure 3E**). These data suggest that following antigenspecific TCR activation, CD8^+^ T cells kill MHC-I-negative tumors via both an antigen-independent and tumor MHC-I-independent mechanism.

### Following TCR Activation, CD8^+^ T Cells Kill MHC-I-Negative Tumors Through a Direct Cell-Cell Contact-Dependent Mechanism

We next examined the mechanistic requirements for killing tumor cells that lack MHC-I. To determine first whether a soluble factor might be involved, we utilized 0.4μm Transwell™plates, which allow cytokines and other soluble mediators, but not cells, to pass through freely. MHC-I-negative CT2A-TRP2-β2mKO tumor cells were placed on the bottom of the Transwell™plates, with the various combinations of TRP-2 T cells and TRP-2 Mφ added to either the same or opposite side of the membrane. Physically separating T cells and/or MΦ from tumor cells obviated tumoricidal activity, suggesting a requirement for direct cell-cell contact rather than a soluble factor (**Figure 4A**).

**Figure 4:**
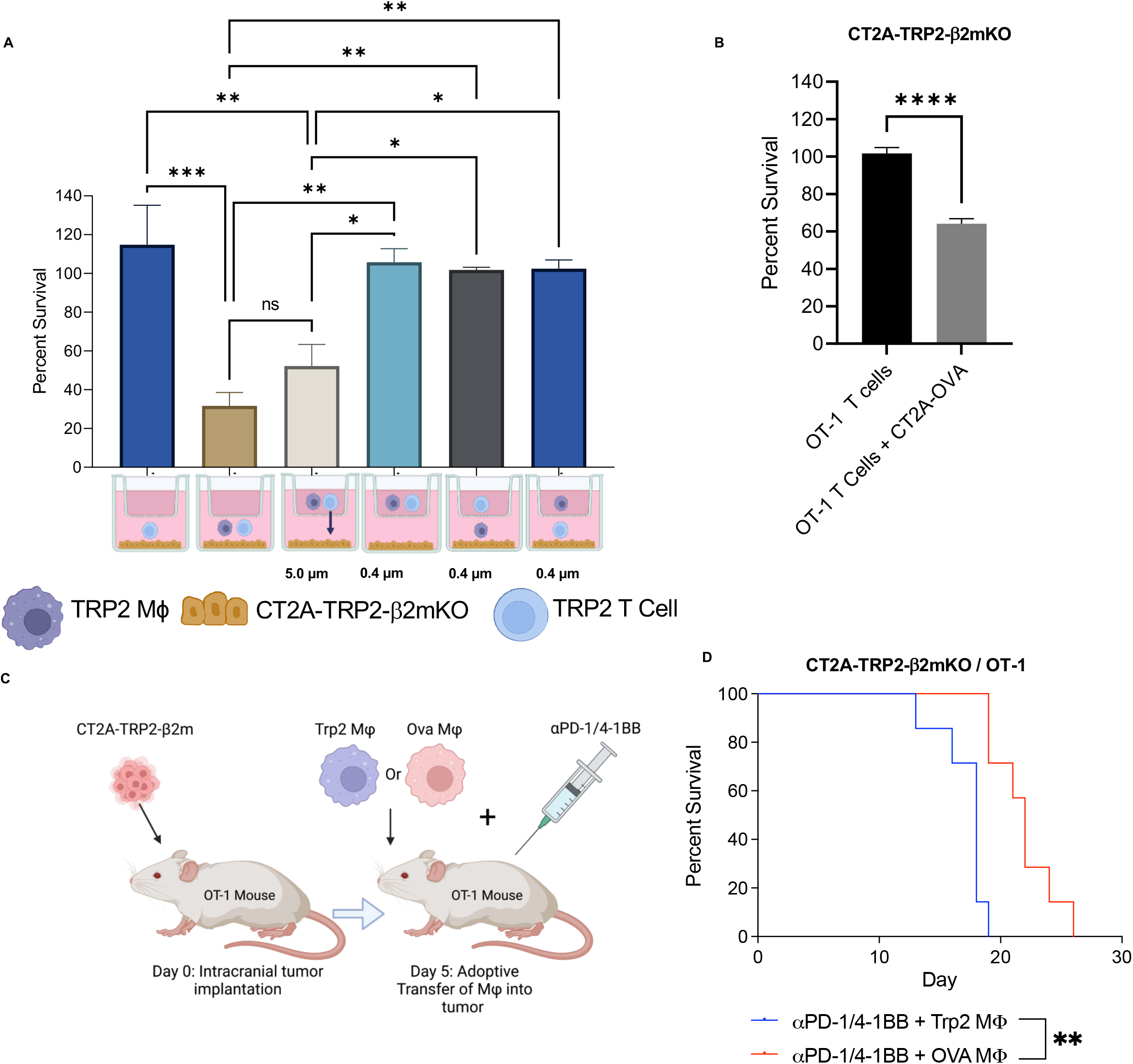
CD8^+^ T cells require contact with APCs and subsequently tumor cells to mediate cytotoxicity against MHC-I-negative tumors. **A**. CT2A-TRP2-β2mKO tumors were co-cultured with TRP2 CD8^+^ T cells and TRP2 Mϕ in different combinations on either side of 5.0μm or 0.4μm Transwell™as depicted. T cells can migrate across 5.0 μm Transwell™inserts whereas macrophages cannot (**Supplementary Fig. 4A).** Neither T cells nor macrophages can migrate across 0.4μm Transwells. After 24 hours of co-culture, the number of remaining viable tumor cells was quantified using flow cytometry and percent survival was calculated in comparison to tumor only wells. Cells were plated at a 10:1 ET. **B**. Percent survival of CT2A-TRP2-β2mKO tumor cells after 24 hours of co-culture with OT-1 T cells either in the presence or absence of OVA antigen stimulation provided by MHC-I-positive CT2A-OVA tumors at a 10:1 ET. Percent survival was calculated as described above. **C**. Schematic of experimental design CT2A-TRP2-β2mKO tumors (**d**) were IC implanted into OT-1 mice. Cognate antigen loaded OVA macrophages (5×10^5^) or non-cognate antigen loaded TRP2 macrophages (5×10^5^) were then IC implanted into the tumor site at day 5 followed by αPD-1/4-1BB immunotherapy. Mice were evaluated for overall survival in days following tumor implantation. N=5 per group. Data in **a** and **b** are shown as mean ±s.e.m. *P* values were determined using one-way ANOVA with post-hoc Tukey’s test for **b**, and by two-tailed, unpaired Student’s t-test for **b**. Data in **a & b** are representative findings of at least 3 technical replicates from one of a minimum of two independently repeated experiments with similar results. Survival in **d** was assessed by Gehan-Breslow-Wilcoxon test. Schematic made with Biorender.com

We then investigated whether Mφ or T cells more directly contributed to the contact-dependent tumor cytotoxicity mechanism. For these experiments, we instead utilized 5.Oμm Transwell™plates, which allow for the smaller T cells, but not the larger Mφ, to pass through to the bottom of the well containing tumor cells (validation in **Supplemental Figure 4A**). CT2A-TRP2-β2mKO tumor cells were placed on the opposite side of the transwell insert from T cells and Mφ. This setup permitted T cells to first make direct contact with Mφ and subsequently pass through the membrane to make direct contact with tumor cells. The extent of tumor killing here was not significantly different than that in which all three cell types were in contact (**Figure 4A**). This suggests that killing of MHC-I-negative tumors requires direct contact between tumor cells and TCR-stimulated T cells, while Mφ need only fill the role of providing antecedent antigen-specific T cell stimulation.

We therefore sought to determine whether it is MΦ exclusively that are required as the source of TCR stimulation, or whether any MHC-I-positive APC might play that role. To assess this, we performed *in vitro* cytotoxicity experiments mixing CD8^+^ T cells with both MHC-I-positive and MHC-I-negative tumor cells, but eliminated MΦ from the culture. This permitted us to determine whether antigen-cognate TCR activation by MHC-I-positive tumor cells might be sufficient to elicit CD8^+^ T cell killing of their MHC-I-negative counterparts. Unsurprisingly, OVA-specific OT-1 T cells cultured only with MHC-I-negative, OVA-negative CT2A-TRP2-β2mKO tumor cells failed to result in tumor killing (**Figure 4B).** However, the simple addition of MHC-I-positive CT2A-OVA tumor cells licensed T cell killing of antigen-mismatched CT2A-TRP2-β2mKO tumor cells (**Figure 4B)**. Interestingly, the addition of antibody blocking LFA-1 and ICAM-1 slightly reduced the *in vitro* cytotoxicity of both MHC-I-positive and -negative tumors, providing further evidence of a T cell-contact-dependent cytotoxicity mechanism (**Supplementary Figure 4B**).

These data suggested that TCR-activated T cells, regardless of the source of antigen presentation, are capable of killing tumor cells lacking both MHC and cognate antigen *in vitro*. We now aimed to test whether such phenomena would persist *in vivo*. Earlier, we demonstrated that aPD-1/4-1BB immunotherapy was ineffective against IC CT2A-TRP2-β2mKO tumors implanted into OT-1 mice (**Figure 2E**). In this model, as all CD8^+^ T cells are specific for OVA, TCR activation does not occur due to the lack of OVA antigen. Our *in vitro* data have suggested that while the killing of MHC-I-negative tumors is tumor antigen-agnostic, cognate antigen is still necessary for antecedent TCR activation. We therefore tested whether injecting cognate antigen-loaded MΦ into the tumor might prove sufficient for licensing an immunotherapeutic response against tumors that lack both MHC-I and the same cognate antigen. We therefore implanted MHC-I-negative, OVA-negative CT2A-TRP2-β2mKO gliomas orthotopically into OT-1 mice and subsequently administered cognate OVA-loaded MΦ or non-cognate TRP2-loaded MΦ into the tumor, followed by treatment with aPD-1/4-1BB immunotherapy (schematic in **Figure 4C)**. OT-1 mice implanted with OVA-MΦ demonstrated a significant survival advantage compared to OT-1 mice implanted with TRP2-MΦ (**Figure 4D)**. These data suggest that *in vivo* TCR activation is both necessary and sufficient to result in the T cell-mediated killing of MHC-I-negative tumors, in tumor antigen-agnostic fashion.

### CD8^+^ T Cell Recognition of MHC-I-Negative Tumors is NKG2D-Mediated in Both Murine and Human Models

In an effort to clarify the nature of the cell-cell contact-dependent mechanism underlying CD8^+^ T cell recognition of MHC-I-negative tumors, we first employed an unbiased approach to examine differences in inflammatory gene expression profiles in antigen-specific CD8^+^ T cells exposed to both MHC-I-negative tumors and cognate antigen loaded MΦ. Gene sets from OVA-activated OT-1 T cells co-cultured with OVA antigen-negative CT2A-TRP2-β2mKO tumor cells or from OVA-activated OT-1 T cells alone were both compared against naïve OT-1 T cells. OVA negative β2mKO tumor lines were used to ensure the absence of tumor-TCR interactions. OT-1 TCR activation for these experiments was accomplished by concomitant culture with OVA MΦ. TCR-activated OT-1s exposed to CT2A-TRP2-β2mKO exhibited increased expression of genes denoting both NK- and T-cell functions, increases not seen simply with TCR-activation of OT-1s (**Figure 5A).** Within the T cell function gene set, differentially expressed genes included activation markers, but not receptors involved in direct contact-mediated cytotoxicity (**Supplementary Figure 4A**). Notably, the expression of TNF family cytotoxic receptor TRAIL, part of the apoptosis gene set, was slightly increased (log_2_ fold-change 0.562, adj. *P* value=0.0262) (**Supplementary Figure 4B**). Therefore, given its known role in contact-dependent T cell cytotoxicity, we investigated whether TRAIL might be involved in killing MHC-I-negative tumors *in vitro*. However, blocking TRAIL did not significantly reduce T cell mediated *in vitro* cytotoxicity directed against either MHC-I-positive or negative tumors (**Supplementary Figure 4C**).

**Figure 5:**
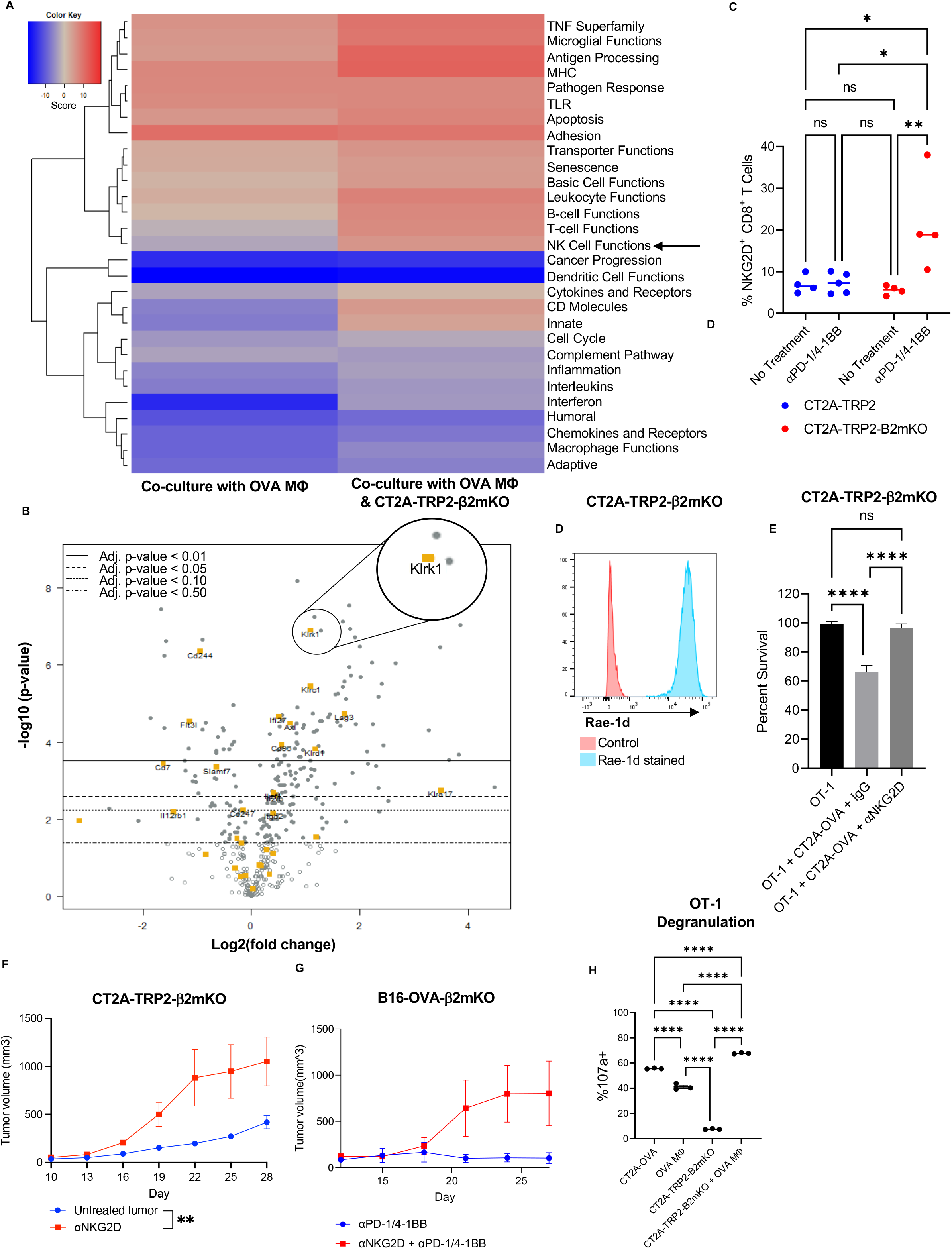
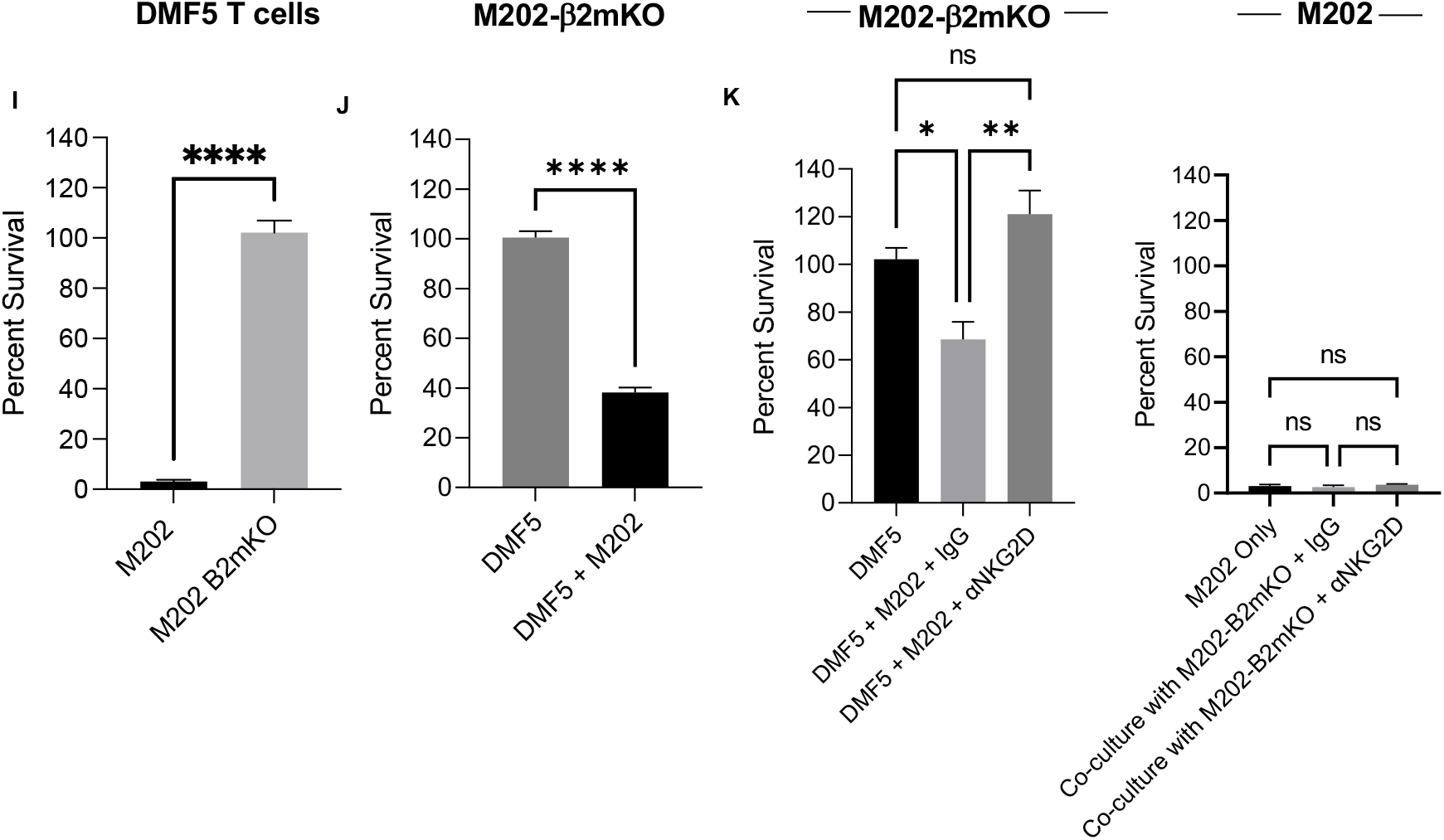
NKG2D on activated CD8^+^ T cells mediates killing of MHC-I-negative tumor cells, but not MHC-I-positive tumors in both Murine and Human tumor models. **A**. Gene set pathway analysis of RNA expression by OT-1 CD8^+^ T cells co-cultured with cognate antigen-loaded OVA Mϕ or OVA Mϕ and CT2A-TRP2-β2mKO, both compared to OT-1 cultured alone in TCM + IL-2. Red indicates increased expression of pathway genes compared to OT-1s cultured alone, blue indicates reduced expression. **B**. Volcano plot showing differential expression of NK cell function genes between OT-1s co-cultured with OVA Mϕ and CT2A-TRP2-β2mKO as compared with OT-1s cultured alone. OT-1s co-cultured with OVA Mϕ and CT2A-TRP2-β2mKO have significantly increased KLRK1 expression (adj. *P* value=3.25×10^-5^), encoding the cytotoxic receptor NKG2D. **C**. Percent NKG2D+ CD8^+^ T cells in tumors of mice bearing CT2A-TRP2-B2mKO or CT2A-TRP2 tumors treated with αPD-1/4-1BB or PBS vehicle. C57BL/6 mice were implanted with CT2A-TRP2 or CT2A-TRP2-β2mKO tumors IC then treated with intraperitoneal αPD-1/4-1BB or PBS at day 18. Tumors were then harvested 24 hours later, dissociated, and assessed by flow cytometry. N= 5/group **D**. NKG2D ligand Rae-1d expression on CT2A-TRP2-β2mKO tumors *in vitro*. **E**. Percent survival of CT2A-TRP2-β2mKO co-cultured with OT-1s alone or in combination with MHC-I-positive-OVA-positive glioma cells (CT2A-OVA) and NKG2D blocking antibody (αNKG2D). **F.** Subcutaneous tumor growth of CT2A-TRP2-β2mKO tumors in C57BL/6 mice treated with either αNKG2D or PBS vehicle control. N=8/group. **G.** Subcutaneous tumor growth curve of B16-OVA-β2mKO, treated with either αPD-1/4-1BB or αNKG2D and αPD-1/4-1BB. **H.** Percent 107a+ OT-1 T cells co-cultured for 6 hrs with OVA Mϕ and MHC-I-positive or negative tumor assessed by flow cytometry (technical triplicate). **I**. Percent survival of MART-1 antigen positive M202 or M202-β2mKO melanoma cells co-cultured with MART-1 specific DMF5 transduced T cells (DMF5). **J** Survival of M202-β2mKO in the presence or absence of MART-1 stimulation provided by MHC-I positive M202 tumor cells **K.** Survival of M202 and M202-β2mKO in co-culture with DMF5 T cells and αNKG2D. Data presented as mean ±s.e.m. unless otherwise stated. Tumor growth curves in **f & g** were compared with twoway repeated measures ANOVA. *P* values were determined using two-way ANOVA with post-hoc Tukey’s test for **c**, oneway ANOVA with post-hoc Tukey’s test for **e,h & k**, and by two-tailed, unpaired Student’s t-test for **i & j**. Cytotoxicity data are representative findings of 6 technical replicates from one of a minimum of two independently repeated experiments with similar results.

Within the NK cell function gene set, the expression of KLRK1, which encodes the cytotoxic receptor NKG2D, was especially prominent in OT-1s exposed to CT2A-TRP2-β2mKO **(Figure 5B**) (log2 fold-change 1.1, adj. *P* value=3.25×10^-5^).NKG2D is an activating receptor traditionally associated with cytotoxic function in NK cells, though it is also expressed on activated CD8^+^ T cells where its role has mainly been described as co-stimulatory [16–19]. In mice, the ligands for NKG2D are the β2m-independent non-classical MHC-I molecules RAE-I, H60, and MULT-1 [16]. Unlike classical MHC-I which is expressed on all nucleated cells, non-classical MHC-I expression is generally stress-induced and is upregulated on tumor cells [18, 20, 21]. The expression of non-classical MHC-I on tumor cells has been shown to be further upregulated following exposure to chemotherapy or ionizing radiation[22].

Similarly, following TCR activation, engagement of the NKG2D receptor by its ligands on a target cell activates cytotoxic effector functions in the NKG2D^+^ cell, including the release of cytotoxic granules and the expression of Fas-L [23–27]. NKG2D effector functions in CD8^+^ T cells, specifically, appear to be dependent on concurrent TCR activation to limit effector activity to appropriate surrounding targets [25, 28]. In murine CD8^+^ T cells, NKG2D expression is upregulated in response to TCR activation (**Supplementary Figure 4D)**. As CD8^+^ T cell-mediated killing of MHC-I-negative tumors in our models was likewise dependent on TCR activation but did not require antigen cognate interactions between T cell and tumor cell, we hypothesized that NKG2D might be an important contributor to the tumoricidal mechanism. To initially assess, we began by examining NKG2D expression on T cells in our models. Accordingly, we looked at differences in NKG2D expression on CD8^+^ T cells infiltrating either CT2A-TRP2 or CT2A-TRP2-β2mKO gliomas in untreated mice or in mice treated with aPD-1/4-1BB. NKG2D levels proved significantly higher on CD8^+^ T cells infiltrating the treated MHC-I-negative CT2A-TRP2-β2mKO tumors when compared to both treated MHC-I-positive tumors and untreated tumors **(Figure 5C).**Detecting such increased levels of NKG2D on T cells, we conversely examined two of our MHC-I-negative tumor lines for the presence of the relevant NKG2D ligands. CT2A-TRP2-β2mKO was found to express the non-canonical MHC I molecules Rae-1 (**Figure 5D)** and MULT-1 (**Supplementary Figure 4E)**, findings that were recapitulated with the additional glioma line GL261-OVA-β2mKO (**Supplementary Figures 4F-G)**.

To examine the functional contribution of NKG2D to CD8^+^ T cell killing of MHC-I-negative tumors, we repeated our prior *in vitro* tumor cytotoxicity studies, this time in the presence or absence of NKG2D blocking antibody (clone CX5). OT-1 T cells were again cultured with OVA-negative, MHC-I-negative CT2A-TRP2-β2mKO tumor cells, although for these experiments, we utilized OVA-positive, MHC-I-positive OVA-expressing tumor cells (CT2A-OVA) as the source of OVA antigen presentation and OT-1 TCR activation. As with our previous results, OT-1 T cells successfully killed CT2A-TRP2-β2mKO tumors, but only when an OVA-presenting cell was likewise present (**Figure 5E)**. Killing of CT2A-TRP2-β2mKO, however, was completely abrogated in the presence of anti-NKG2D blocking antibody (**Figure 5E**). The use of CT2A-OVA tumor as a source of OVA TCR stimulation also permitted us to evaluate the impact of anti-NKG2D on concomitant killing of MHC-I-positive tumor cells. No such impact was seen, as CT2A-OVA was readily killed under either condition (**Supplementary Figure 4H)**.

To determine whether blocking NKG2D might similarly abrogate the killing of MHC-I-negative tumors *in vivo*, we subcutaneously implanted CT2A-TRP2-β2mKO tumors into C57BL/6 mice and treated with αNKG2D or vehicle control. Mice treated with αNKG2D exhibited a significantly increased rate of tumor growth, akin to what was seen in our *in vitro* models (**Figure 5F)**. To additionally assess the impact of NKG2D blockade on a more potent mode of tumor kill involving immunotherapy, we utilized ICB-responsive B16 melanomas treated concomitantly with aPD-1/4-1BB. Administering αNKG2D reduced the efficacy of aPD-1/4-1BB in these tumors, increasing the rate of tumor growth (**Figure 5G).**

The downstream cytotoxic effectors of NKG2D signaling can include granzymes, perforin, and Fas-L [27, 29], raising the question as to which of these might be mediating MHC-I-negative tumor cell kill in our system. Our expression analysis data revealed an increase in Granzyme B and Fas-L expression (**Supplementary Figures 4I,J**) in TCR-activated T cells exposed to MHC-I-negative tumors, suggesting degranulation may play a role in the tumoricidal mechanism. Indeed, degranulation by OT-1 T cells was increased in the presence of MHC-I negative tumor and cognate antigen loaded macrophages as compared with macrophages alone (**Figure 5H**). Given the results of prior studies [30–32], we additionally specifically tested the role of Fas-L/Fas interactions. Fas-L expression on tumor infiltrating CD8^+^ T cells was increased in response to immunotherapy, but was not significantly different across CD8^+^ T cells isolated from MHC-I-positive vs MHC-I-negative tumors (**Supplementary Figure 4K**). Additionally, *in vitro*, antigen-negative Fas receptor knockout (FasKO) melanomas were still susceptible to killing by TCR-activated CD8^+^ T cells (**Supplementary Figure 4L**), suggesting Fas-L/Fas interactions are not critical for the tumoricidal mechanism.

Having examined the role of NKG2D both *in vitro* and *in vivo* in mice, the question remained as to whether similar mechanisms for killing of MHC-I-negative tumors might be at play in human cancers. To address the translational relevance, we performed *in vitro* cytotoxicity experiments using a MART-1-expressing M202 melanoma line (HLA-A*0201) or an MHC-I-negative β2mKO version of the same cells (M202-β2mKO). Tumors were co-cultured with DMF5 TCR-transduced human T cells (DMF5 T cells), which recognize MART-1 presented in the context of HLA-A*0201 [33–35]. Similar to seen in our murine experiments, DMF5 T cells alone efficiently killed MHC-I-positive M202 cells, but failed to kill M202-β2mKO, unless undergoing prior MART-I TCR stimulation **(Figure 5I, 5J)**. Sham-transduced human CD8^+^ T cells failed to kill either M202 or M202-β2mKO cells in vitro (**Supplementary Figure 4M**). Furthermore, the addition of aNKG2D entirely abrogated the killing of MHC-I-negative M202-β2mKO but had no effect on the killing efficiency of MHC-I-positive M202 (**Figure 5K)**. These data suggest that NKG2D-mediated T cell killing of MHC-I-negative tumors is pertinent to and persists amidst human cancer.

## Discussion

Traditional models of anti-tumor immunity have centered on tumor cell killing as a function of CD8^+^ T cell recognition of cognate antigen presented exclusively in the context of tumor cell MHC-I molecules [1, 2, 9, 36]. Here, we report a novel T cell mechanism for tumor cytotoxicity that depends neither on tumor antigen, nor on tumor MHC-I expression. This mechanism is particularly active and effective against MHC-I-loss variants and is mediated instead by T cell NKG2D engagement of non-classical MHC-I (NKG2D ligands) on tumor cells in antigen-independent fashion. Subsequent tumor cell kill depends on prior TCR-mediated T cell activation (even if by ultimately tumor-irrelevant antigen) and appears to involve granzyme and/or perforin. The mechanism remains active in tumors containing a mixture of MHC-I-replete and -deficient cells, where TCR activation can seemingly be provided by either MHC-I-replete tumor cells or adjacent myeloid APCs. Likewise, the mechanism extrapolates to human tumors and T cells, suggesting clinically relevant findings.

Our findings call into question the conventional notion that tumor downregulation of MHC-I necessarily represents an effective mode of immune escape [37–39]. Results from recent clinical studies appear to corroborate that such notions may indeed be faulty [11, 40–44]. Within GBM alone, for instance, two separate studies gathering single cell RNA sequencing data from human tumors have found β2m expression to be inversely correlated with survival [11, 44]. Likewise, additional studies have implied that MHC-I loss does not necessarily render tumor cells invulnerable to CD8^+^ T-cell dependent immunotherapies, including ICB [10, 45]. The perception of MHC-I loss as a means of cancer immune-escape, then, is perhaps best described as controversial.

While some studies to date have indeed highlighted a possible prognostic benefit to tumor MHC-I loss, these have often attributed such benefit to a sudden conferred susceptibility to NK cell-mediated killing [10, 40, 43]. Strikingly, however, we demonstrate that NK cells are not required for the efficacy of an ICB-involving regimen employed against MHC-I-negative tumors. In contrast, we show the requirement for CD8^+^ T cells is indeed maintained for immunotherapeutic efficacy, despite the absence of tumor MHC-I to facilitate a traditional T cell killing mechanism.

An additional interesting facet of our data is the apparent antigen-independence of the observed T cell killing mechanism. While CD8^+^ T cell cytotoxicity in our studies required prior TCR activation, subsequent TCR-mediated recognition of tumor cells was not required. Therefore, tumors cells lacking MHC-I needed not present or even express the same antigen that was enlisted for T cell activation. Instead, only tumor expression of NKG2D ligand was required, and tumor cells lacking MHC-I appeared to have a reflexive increase in these ligands, leaving them more subject to this mode of cell kill. The implication, however, is that T cells specific for irrelevant antigens need only be activated in the vicinity of tumors lacking MHC-I, but possessing NKG2D ligands, to affect a cytotoxic response. The requirement for NKG2D ligands on target cells, then, would be the only restriction limiting otherwise non-antigen restrained T cell activity. Another implication is that even polyclonally or agnostically activated T cells can become potential effectors in this cytotoxicity model.

These findings, while surprising, are perhaps not entirely without precedent. While the role of NKG2D in CD8^+^ T cells has mainly been described as co-stimulatory, ([16–19], it has previously been shown that NKG2D receptor-ligand interactions can lead to the formation of immunologic synapses between T cells and target cells in an antigen-independent manner [17]. Our findings of a contact-dependent cytotoxicity mechanism with evidence of degranulation support this notion of immunologic synapse formation by NKG2D in an antigen-independent manner.

Interestingly, Upadhyay et al. recently described a bystander killing phenomenon by chimeric antigen receptor (CAR) T-cells in the context of mixed antigen-positive and antigen-negative tumors [30]. While CAR-T-cells act in an MHC-independent fashion regardless, the authors also reported the ability of endogenous bystander T cells to kill a mixed MHC-I-positive and negative tumor *in vitro*, in seemingly Fas-dependent fashion. Direct activation of the TCR was required and was provided by MHC-replete tumor cells. While we uncovered a similar phenomenon, we also show that TCR stimulation can come from non-tumor sources (i.e. myeloid cells) and is thus sufficient to mediate killing of tumors entirely devoid of MHC-I. Our findings are further contrasted as we additionally highlight a novel role for NKG2D in mediating tumor cell recognition and kill, a mechanism that persists when Fas is absent. We instead suggest a role for perforin and granzyme in mediating tumor cell kill following NKG2D-based tumor recognition.

To our knowledge, then, this remains the first characterization of an MHC-I- and antigen-independent CD8^+^ T-cell mechanism for tumor cell kill that is seen consistently *in vivo*, as well as in human tumor cells. It is also the first to clearly demonstrate that T cell-dependent immunotherapies can indeed remain effective against tumors, even when uniformly lacking MHC-I. Further investigations into the role of NKG2D and NKG2D ligands in mediating tumor immunotherapeutic susceptibility, as well as into other potential therapeutic implications for these findings, are warranted.

## Author contributions

EL and KW conceived and designed the experiments. EL, VDA, XC, DW, JWP, KW acquired the data. EL, WT, PEF, and KW analyzed and interpreted the data. EL and KW wrote the manuscript. All authors: read, revised, and approved the final manuscript.

## Acknowledgements

We would like to thank A. Ribas for providing the M202 and M202 B2mKO cell lines used in this manuscript, as well as K. Wood for providing the Yummer FasKO cell line. The work was supported in part by the Cancer Research Institute Lloyd J. Old STAR Award (P.E. Fecci) and the Duke Office of Physician Scientist Development (E. Lerner, K. Woroniecka).

## Materials and Methods

### Mice

The Institutional Animal Care and Use Committee (IACUC) approved all experimental procedures. Animal experiments involved the use of female mice at 6-12 weeks of age. C57BL/6 mice were purchased from Charles River Laboratories. OT-1 (#003831), and CD8KO (#002665) were purchased from Jackson Laboratories. Mice were housed at the Duke University Medical Center Cancer Center Isolation Facility (CCIF) under pathogen-free conditions.

### Cell lines

Murine cell lines included CT2A glioma, GL261 glioma, B16 melanoma, and Yummer-FasKO melanoma. The YUMMER-FasKO cell lines were kindly provided by Dr. Kris Wood (Duke University). All murine cell lines are syngeneic in C57BL/6 mice. We transfected CT2A to express the antigen Trp2 to generate CT2A-Trp2. Additionally, we transfected GL261 and B16 to express OVA to generate GL261-OVA and B16-OVA, respectively. To knockout β2m in CT2A-Trp2, GL261-OVA and B16-OVA, we utilized a CRISPR-based strategy using LentiCRISPRv2 plasmid (#52961, Addgene) as described previously[46]. Briefly, the lentiCRISPRv2 vector was digested with BsmBI (NEB), gel purified using the QIAquick Gel Extraction Kit (Qiagen), ligated with phosphorylated oligonucleotides, and transformed into Stbl3 bacteria. Positives clones were verified and sequenced. We used gRNA that targets the 1st exon of the β2m gene and the sequence is as follows: 5’ CCGAGCCCAAGACCGTCTAC. HEK 293T-cells were then transfected with LentiCRISPRv2-β2m, psPAX2 (Addgene #12260), and pVSVg (Addgene #8454). Transfected cells were cultured for 48h, then supernatant was collected and tumor cells were transduced with Polybrene Infection/Transfection Reagent (Sigma-Aldrich) at 8ug/mL. Cells were cultured in viral supernatant for 24h. Media was changed and cells were cultured for another 5 days. H-2Kb and H-2Db negative cells were single-cell sorted and confirmed by flow cytometry. The human melanoma M202 and M202-β2MKO cell lines were kindly provided by Dr. Antoni Ribas (UCLA). All cell lines with the exception of YUMMER were cultured *in vitro* in Dulbecco’s Modified Eagle Medium (DMEM) with 2mM l-glutamine and 4.5 mg/mL glucose (Gibco) containing 10% fetal bovine serum (FBS). YUMMER cells were cultured in DMEM/F12 containing 10% FBS and supplemented with non-essential amino acids.

### In vivo survival experiments

For intracranial implantation, tumor cells were mixed 1:1 with 3% methylcellulose and loaded into a 250 μl syringe (Hamilton). The needle was positioned 2 mm right of the bregma and 4 mm beneath the surface of the skull using a stereotactic frame. Cells were administered in 5 μl of volume. For CT2A-TRP2 2.5 x 10^4^ cells were injected, for CT2A-TRP2-β2mKO 5.0 x 10^4^ cells were injected. These doses were determined by tumorgenicity experiments to determine dose required for a median survival of around 21 days.

For subcutaneous implantation, the appropriate number of tumor cells were administered in 200 μl PBS under the skin of right flank. For B16-OVA-β2mKO 2.5 x 10^5^ cells were injected subcutaneously. For CT2A-TRP2-β2mKO, 1 x 10^6^ cells were injected subcutaneously. Tumors were measured every 3 days. Mice were sacrificed when tumor volume reached 2000mm^3^, tumors were over 20mm in one direction, or tumors became ulcerated.

Tumors were implanted into the indicated mouse strain for each experiment. In adoptive transfer experiments with TRP-2-specific T-cells, we administered 10^7^ TRP-2 TCR-transduced T-cells intravenously (iv) 7 days post-tumor implantation. For adoptive transfer experiments with antigen-loaded macrophages, we pulsed macrophages with OVA 257-264 (SIINFEKL) peptide and then administered 5 x 10^5^ macrophages intracranially at the tumor site 5 days post-tumor implantation. Unless otherwise indicated, mice were monitored for survival or sacrificed once they reached a humane endpoint.

### *In vivo* antibody treatment

Antibodies for *in vivo* treatment were obtained from Bio X Cell. These include anti-mouse PD-1 (RMP1-14, Catalog #BE0146), anti-mouse 4-1BB (LOB12.3, Catalog #BE0169), anti-mouse CD8α (2.43, Catalog #BE0061), anti-mouse CD4 (GK1.5, Catalog #BE0003-1), anti-mouse NK1.1 (PK136, Catalog #BE0036), anti-mouse NKG2D (HMG2D, Catalog # BE0111). For *in vivo* treatment with aPD-1 and a4-1BB antibody, 200 μg each of PD-1 and 4-1BB was diluted with PBS for a total injection volume of 200 μL. Mice received intraperitoneal (IP) injections of immune checkpoint blockade (PD-1 and 4-1BB) every 3 days, starting on day 9 after tumor implantation, for a total of 4 treatments. unless otherwise specified. For antibody depletions, 200 μg of each antibody was diluted with PBS for a total injection volume of 200 μL. For depletions beginning prior to tumor implantations, mice received IP injections each day for 3 days, with the last dose given one day prior to tumor implantation. Maintenance doses were then given every 6 days. Control treatments consisted of 200 μL PBS. Depletion of CD8 (Biolegend APC anti-mouse CD8a clone QA17A07, cat. 155005), NK (Biolegend AF488 NK1.1 clone PK136 cat.108718), (Biolegend PE NKp46 cat. 137611) cells, and CD4 (Biolegend PE CD4 Clone GK1.5, cat. 100408) cells were validated by flow cytometry of blood samples prior to tumor implantation and again in blood of tumor bearing mice one day prior to the scheduled maintenance dose. For vivo blockade of NKG2D, 250ug of aNKG2D was given starting day 8, continuing every 3 days until humane endpoints were reached[47].

### Generation of Antigen-Specific T Cells

We engineered TRP-2 TCR T-cells by retroviral transduction of T-cells with the pMX-TRP-2-TCRβ-2A-α vector (a kind gift Dr. Schumacher, Netherlands Cancer Instute) as described previously [48]. This TCR recognizes the TRP-2_180-188_ epitope in the context of H-2Kb. Briefly, we transfected HEK293T-cells with vectors encoding the TRP-2 TCR and retroviral packaging genes to generate TCR encoding retrovirus. TRP-2 TCR retrovirus was then used to transduce mouse T-cells 48 h after activation with Concanavalin A (ConA) in the presence of 50 U/mL human IL-2. Transduction was performed on non-tissue culture 24-well plates previously coated with 0.5mL of RetroNectin (Takara) at a concentration of 25 μg/mL in PBS. Cells were split every 48h for 5 days in T-cell media (RPMI with 10% FBS, non-essential amino acids, l-glutamine, sodium pyruvate, beta-mercaptoethanol, pen/strep, and gentamycin) prior to their use.

We engineered human T-cells to express the DMF5 TCR that recognizes the MART-1 antigen expressed by the human melanoma M202 cell line [8, 33, 34]. DMF5 TCRα and TCRβ were cloned into the MSGV1 retroviral backbone as a bicistronic message with a P2A self-cleaving peptide separating the α and β domains. DMF5 TCR retrovirus was generated by transfecting HEK293T cells with the DMF5 TCR MSGV1 vector and separate vectors encoding Gag-Pol and Gibbon-ape leukemia virus (GALV) envelope. Anonymous healthy donor peripheral blood mononuclear cells (PBMCs; StemCell) were activated with anti-CD3 (OKT3; Biolegend) in the presence of 100 U/mL human IL-2. After 48 h, activated T-cells were transduced with DMF5 TCR retrovirus to generate DMF5 T-cells. DMF5 T-cells were split into fresh T-cell media (RPMI supplemented with 10% FBS, l-glutamine, HEPES, sodium pyruvate, non-essential amino acids, and pen/strep) plus 100 U/mL IL-2 every 48 h.

OT-1 T-cells were isolated from OT-1 mice by culturing OT-1 splenocytes in T-cell media supplemented with 50IU/mL IL-2 and 1 μM OVA SIINFEKL peptide (Anaspec) for 48 h. Cells were purified for CD8 T-cells as described above and subsequently cultured in TCM with 50IU/mL IL-2, splitting every 24h for a total of 4 days.

### Monocyte purification, macrophage differentiation, and antigen loading

Ly6C^hl^ monocytes were purified from bone marrow (BM) as previously described [49, 50]. Briefly, BM was flushed from the tibiae, femora, humeri, and sternum of C57BL/6, CD45.1 (B6.SJL-Ptprc^a^ Pepc^b^/BoyJ) mice or B2mKO mice (B6.129P2-B2mKO^tm1Unc^ / DcrJ, Jackson Laboratory stock 002087) into cRPMI-10 medium (glutamine-free RPMI-1640 medium with 10% fetal bovine serum, 100U/mL penicillin, 100 μg/mL streptomycin, 100 μM MEM-nonessential amino acids, 2 mM L-glutamine, and 1 mM sodium pyruvate). Red blood cells were lysed with ammonium-chloride-potassium buffer at room temperature for 2 minutes and neutralized with cRPMI-10. The cell suspension was then passed through a 70-μm nylon cell strainer and incubated for 30 minutes at 4 °C in separation buffer (0.5 % bovine serum albumin, 2 mM EDTA in PBS) containing biotinylated anti-CD3ε, anti-CD4, anti-CD8α, anti-CD11c, anti-CD19, anti-B220, anti-CD49b, anti-I-A^b^, anti Sca-1, anti-c-Kit, anti-TER-119, and FITC-conjugated anti-Ly6G and anti-CCR3 (5 μL/mL for anti-CCR3; all others at 1.25 μl/mL). Cells were then washed with labeling buffer (2 mM EDTA in PBS), then incubated for 15 min at 4 °C in labeling buffer containing streptavidin-conjugated and anti-FITC MicroBeads (Miltenyi Biotec). Cells were negatively selected using MACS LD columns per the manufacturer’s instructions. The resulting classical monocytes (>90% purity) were cultured for 5-7 days in non-tissue culture treated dishes containing macrophage differentiation media (glutamine-free RPMI-1640 medium with 20% fetal bovine serum, 100U/mL penicillin, 100 μg/mL streptomycin, 100 μM MEM-nonessential amino acids, 2 mM L-glutamine, 1 mM sodium pyruvate, and 50 ng/mL recombinant murine M-CSF (Peprotech 315-02). Monocyte derived macrophages were loaded with antigen by adding 10 μM peptide dissolved in DMSO to the culture for 4 h, or equal volume of DMSO vehicle only control. Bone marrow derived macrophages (BMDMs) were then washed twice with PBS to remove remaining antigen, separated from culture plates using non-enzymatic dissociation agent (Corning Cellstripper Product # 25-056-Cl) and counted.

### Tissue Processing and Flow Cytometry

A detailed protocol for processing and staining tumors can be found here[51]. In brief, after perfusion with PBS + heparin 1%, tumor-bearing hemispheres were harvested. A single cell suspension was generated and passed through a 70-μm filter. After RBC Lysis (RBC Lysis Buffer, Thermo Fisher Scientific) for 3 min, myelin was removed from the sample with Percoll (Sigma Aldrich) centrifugation. Cells were then resuspended at 1 × 10^6^ ml^-1^ in 100 μl PBS and transferred to a 96-well plate. Before further staining, samples were resuspended in Zombie Aqua Viability Dye (1:400, Biolegend) and incubated for 30 min on ice.

For extracellular staining of harvested or *in vitro* cultured cells, samples were incubated with Fc blocking solution (1:100 anti-mouse CD16/32, Biolegend Catalogue # 101302) in FACS buffer (1x PBS with 2% fetal bovine serum). After blocking, samples were incubated with antibodies (Supplementary Table 1) for 30 min on ice. Stained samples were then fixed in 2% formaldehyde in PBS on ice for 15 min.

Before acquisition, 10 μl of Accucheck Counting Beads (Thermo Fisher Scientific) were added to each sample. To calculate the number of cells per gram of tumor the following calculation was used: number of acquired cells × (number of input beads/number of acquired beads) × (1 / fraction of sample stained) × (1 / tumor weight). Samples were acquired on a LSRFortessa (BD Biosciences) using FACS Diva software v.9 (BD Biosciences) and analyzed using FlowJo v.10 (BD Biosciences).

Mouse-specific antibodies were purchased from BD Biosciences, eBioscience, R and D systems or Biolegend. Tissue processing and flow cytometry was performed as described previously [52] For CD107a staining, OT-1 T cells were co-cultured with OVA loaded BMDMs, CT2A-OVA tumor, CT2A-TRP2-B2mKO tumor, or alone in TCM supplemented with IL-2. Macrophages were stained with CellTrace™ Violet. At the start of co-culture, anti-mouse CD107a antibody (Supplementary Table 1) was added, along with Golgistop (2μM; BD Biosciences) to prevent antibody breakdown. After 5 hours of co-culture, the cells were Fc blocked then stained for CD8 (BV650, table 1), washed, and formalin fixed [53].

### In vitro cytotoxicity assays

In vitro cytotoxicity assays were performed by co-culturing sorted CD8 T-cells (Miltenyi Biotec catalogue # 130-104-075) with tumor cells in the presence or absence of antigen-pulsed macrophages. Prior to co-culture, target-cells were labelled with CellTrace Violet (CTV) (Invitrogen) or CellTrace Far Red (CTFR) and macrophages were labelled with CellTrace CFSE per manufacturer’s protocol to distinguish each cell type. Trp2 T-cells, OT-1 T-cells, DMF5 T-cells, or sham transduced T-cells were co-cultured with the designated target-cell in T-cell media for 24 h in 96-well flat-bottom plates. After 24 h, cells were detached using trypsin and resuspended in 100uL FACS with 10ul CountBright™ beads (Life Technologies Absolute Counting Beads Catalogue # C36950) per well. Remaining viable tumor cells were quantified with flow cytometry. Percent lysis was calculated by counting remaining viable tumor cells in experimental wells versus tumor only control wells, normalized as cells per bead and expressed as percent survival compared with tumor only control wells (% survival = (experimental well viable cells/bead count / tumor only well viable cells/ bead count)*100. In vitro cytotoxicity assays were performed with the following tumor cell lines: CT2A-β2mKO, CT2A-TRP2-β2mKO, CT2A-TRP2, GL261-OVA, GL261-OVA-β2mKO, YUMMER FasKO, M202, and M202-β2mKO.

For cytotoxicity assays with blocking antibodies, 10 μg ml^-1^ anti-mouse NKG2D (Clone HMG2D, BioXCell Catalog # BE0111), anti-mouse TRAIL (Clone N2B2, Invitrogen Catalog #16-5951-85),anti-mouse ICAM-1 (Clone YN1/1.7.4, BioLegend Catalogue #116101), anti-mouse aLFA-1 (Clone M17/4, Biolegend Catalogue # 101118), or anti-human NKG2D (Clone 1D11, BioXCell Catalogue #BE0351) were added 30 to T cells 30 minutes prior to coculture with tumor cells. Remaining viable tumor cells were quantified after 24h as described above.

### In vitro Transwell Cytotoxicity Assays

Transwell in vitro cytotoxicity experiments were performed as described above using 6.5mm 0.4 μm and 5.0 μm Transwell^®^ inserts (Corning Costar 0.4 μm product number 3470, 5.0 μm product number 3421) with the following modifications. CT2A-OVA-β2mKO and GL261-Ova-β2mKO tumor cells were counted, stained with CellTrace™ Violet, and plated on the bottom of 24 well plates. Bone marrow derived macrophages (BMDMs) were loaded with TRP-2 peptide as described above and then stained with CFSE. TRP-2 macrophages and TRP-2-specfic T-cells were then plated in the basket of the Transwell at a 10:1 ratio of T-cells to tumor and 5:1 ratio of macrophages to tumor. After 24 h of culturing, wells were dislodged and quantified by flow cytometry. The inability for macrophages to pass through the 5.0 uM Transwell barrier was confirmed by the absence of CFSE positive cells.

### RNA expression Analysis

OT-1 T-cells were co-cultured either alone in T-cell media supplemented with IL-2 (50IU/mL IL-2), with OVA loaded macrophages, or with both OVA loaded macrophages and CT2A-TRP2-β2mKO tumor cells. Cells were cultured at a 5:1 T-cell to tumor ratio, and 2:1 T-cell to macrophage ratio. After 24 h of co-culture, CD8 T-cells were FACS sorted, and RNA extracted (RNeasy Mini Kit, Qiagen). RNA was analyzed on an nCounter MAX Analysis System (Nanostring) with the PanCancer Immune profiling panel (Nanostring) according to manufacturer instructions. Expression data were analyzed using nSolver and nSolver Advanced Anaylsis software.

### Statistical Analysis

Statistical analysis was conducted in GraphPad Prism version 5.0 (GraphPad Software, La Jolla, CA), primarily using two-tailed, unpaired t-tests or one-way ANOVAs to compare means across groups with a designated significance level of 0.05. Analyses were adjusted for multiple comparisons using the Bonferoni adjustment as indicated. Bar graphs and dot plots display the mean +/- the standard error of the mean. Kaplan-Meier curves were generated for survival analyses and the Gehan-Breslow-Wilcoxon test was used to compare curves. The statistical tests employed for each data presentation are designated in respective figure legends

---

**Supplementary Figure 1.**
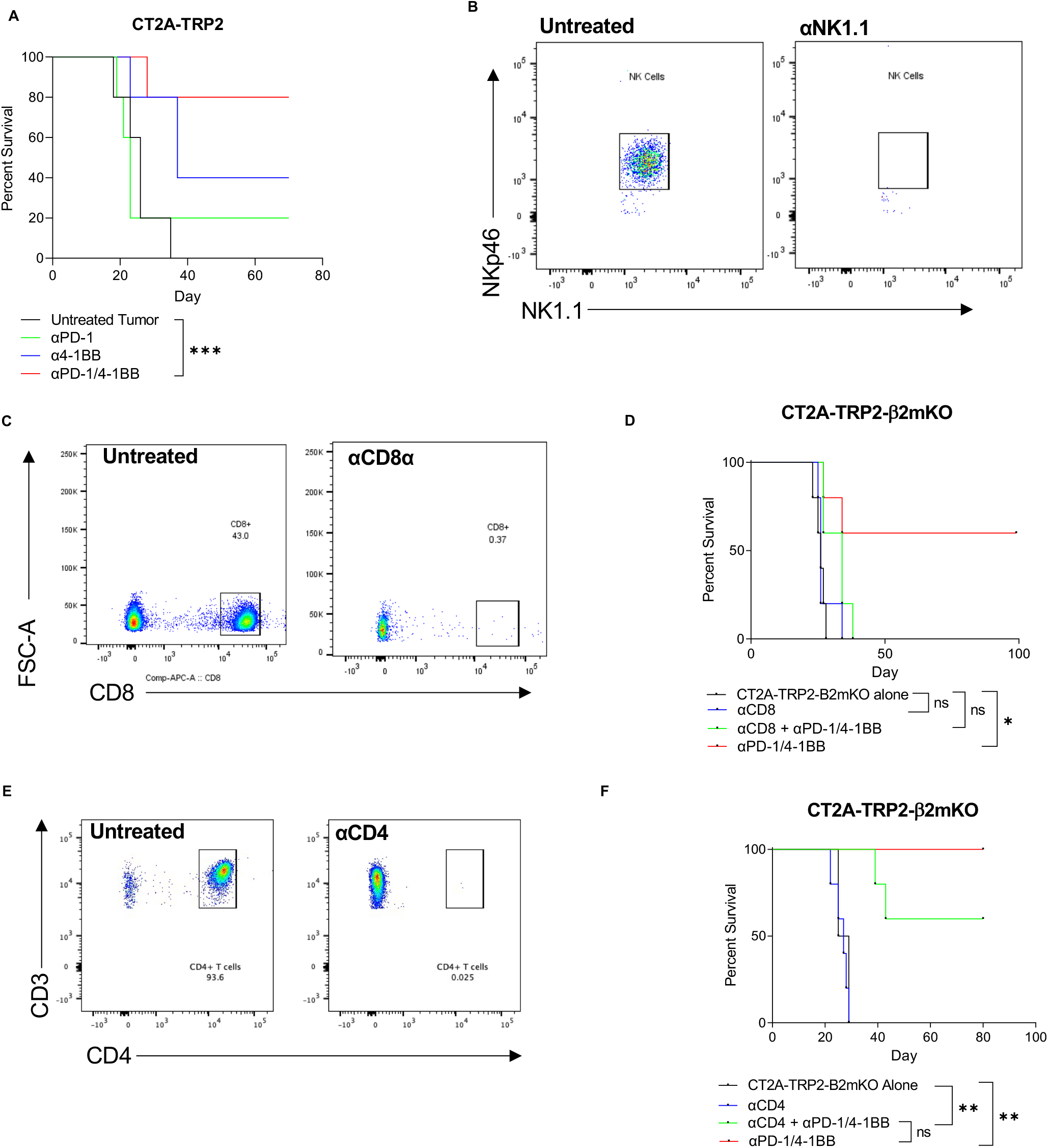
**A.** Kaplan-Meier survival curve of mice implanted with CT2A-TRP2 IC then treated with αPD-1, α4-1BB, αPD-1/4-1BB, or PBS vehicle control. N=5/group. **B**. *In vivo* depletion validation of NK cells from blood of tumor bearing mouse. Depletion >99% by count. **C.** *In vivo* depletion validation of CD8^+^ T cells from blood of tumor bearing mouse. Depletion >99% by count. **D**. Kaplan-Meier survival curve of mice implanted with CT2A-TRP2-B2mKO IC then treated with αPD-1/4-1BB in the presence or absence of CD8 depletion with αCD8α. **E**. *In vivo* depletion validation of CD4^+^T cells from blood of tumor bearing mice. Depletion >99% by count. **F**. Mice were implanted with CT2A-TRP2-B2mKO IC then treated with αPD-1/4-1BB in the presence or absence of CD4^+^ T cell depletion with αCD4. Antibody depletion for **d** and **f** was started three days prior to tumor implantation, with maintenance depletion continued for the duration of the experiment. N=5/group. Survival in **a,d&f** was assessed by Gehan-Breslow-Wilcoxon test. *P* values were Bonferroni corrected for multiple comparisons.

**Supplementary Figure 2.**
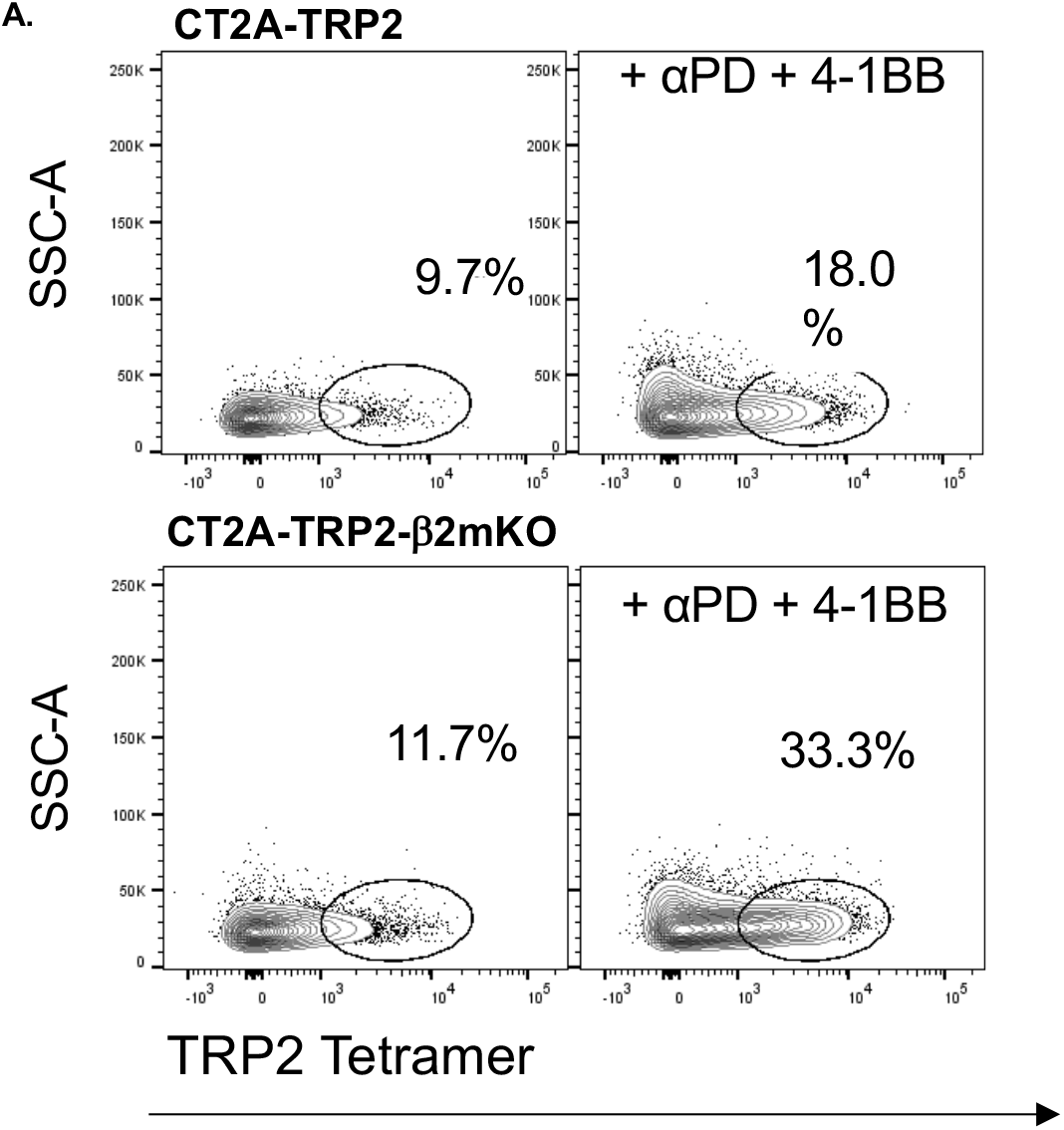
TRP2 Tetramer staining of CD8^+^T cells isolated from IC CT2A-TRP2 or CT2A-TRP2-β2mKO tumors.

**Supplementary Figure 3:**
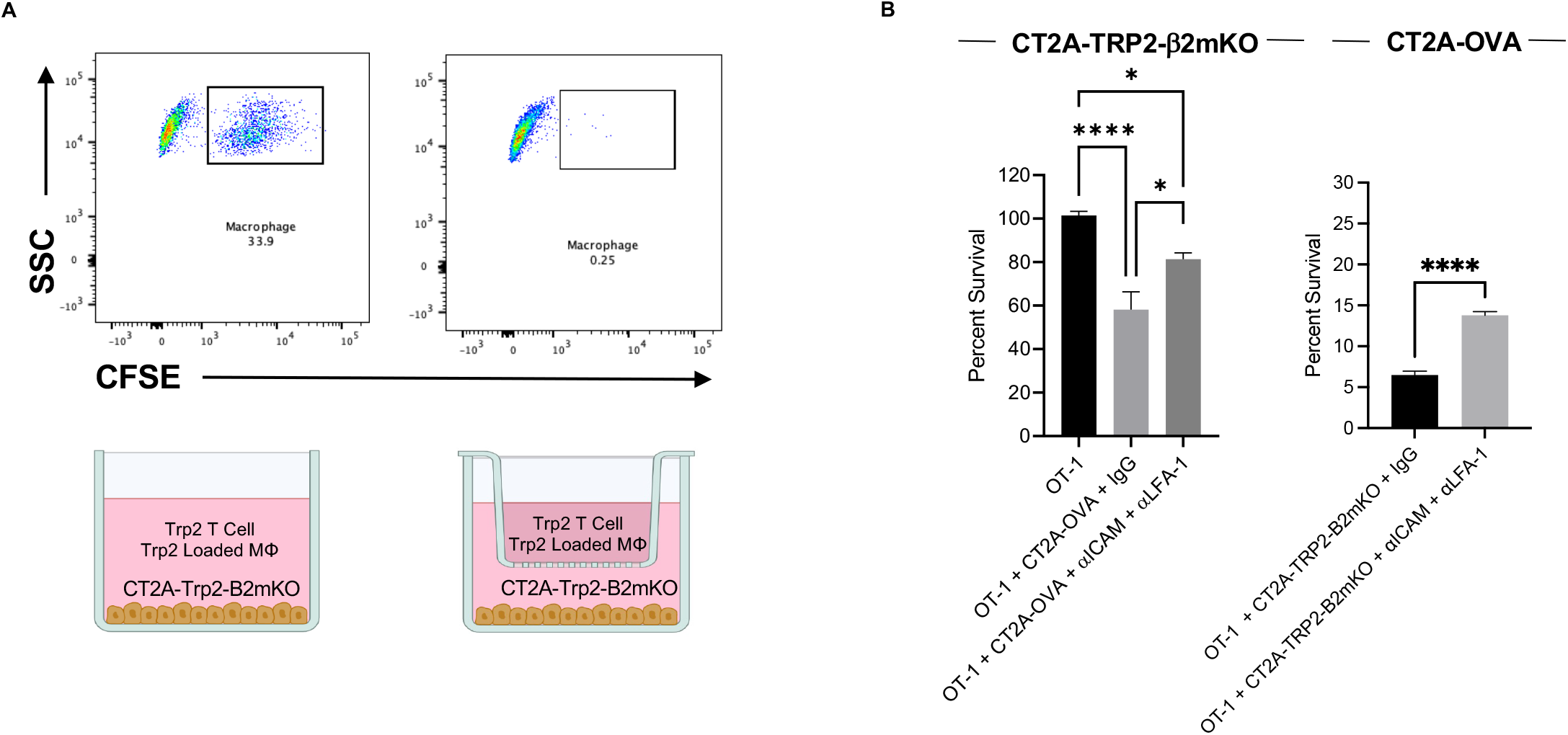
**A**. CFSE-stained acrophages do not pass through 5.0μm transwell membranes. TRP2 T cells, TRP2 Mϕ and CT2A-TRP2-B2mKO tumors were co-cultured either together or separated by a 5.0μm transwell insert, as depicted. After 24 hours of co-culture, the transwell inserts were removed and the remaining cells were assessed with flow cytometry. **B.** Percent survival of CT2A-TRP2-β2mKO or CT2A-TRP2 co-cultured with OT-1s alone, or combination with MHC-I-positive OVA positive glioma cells (CT2A-OVA) and blocking antibody for ICAM-1 and LFA-1 or IgG isotype control.

**Supplementary Figure 4:**
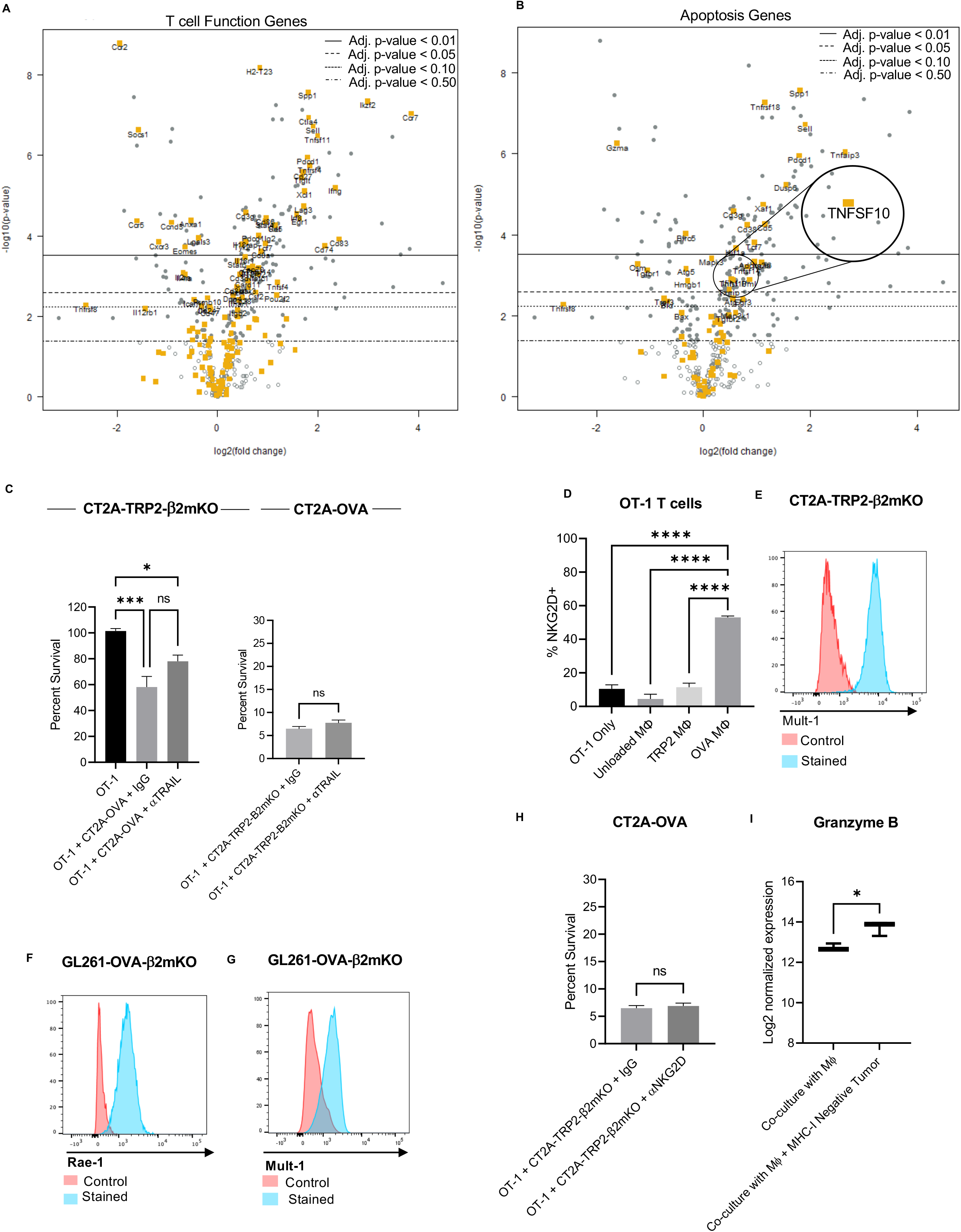
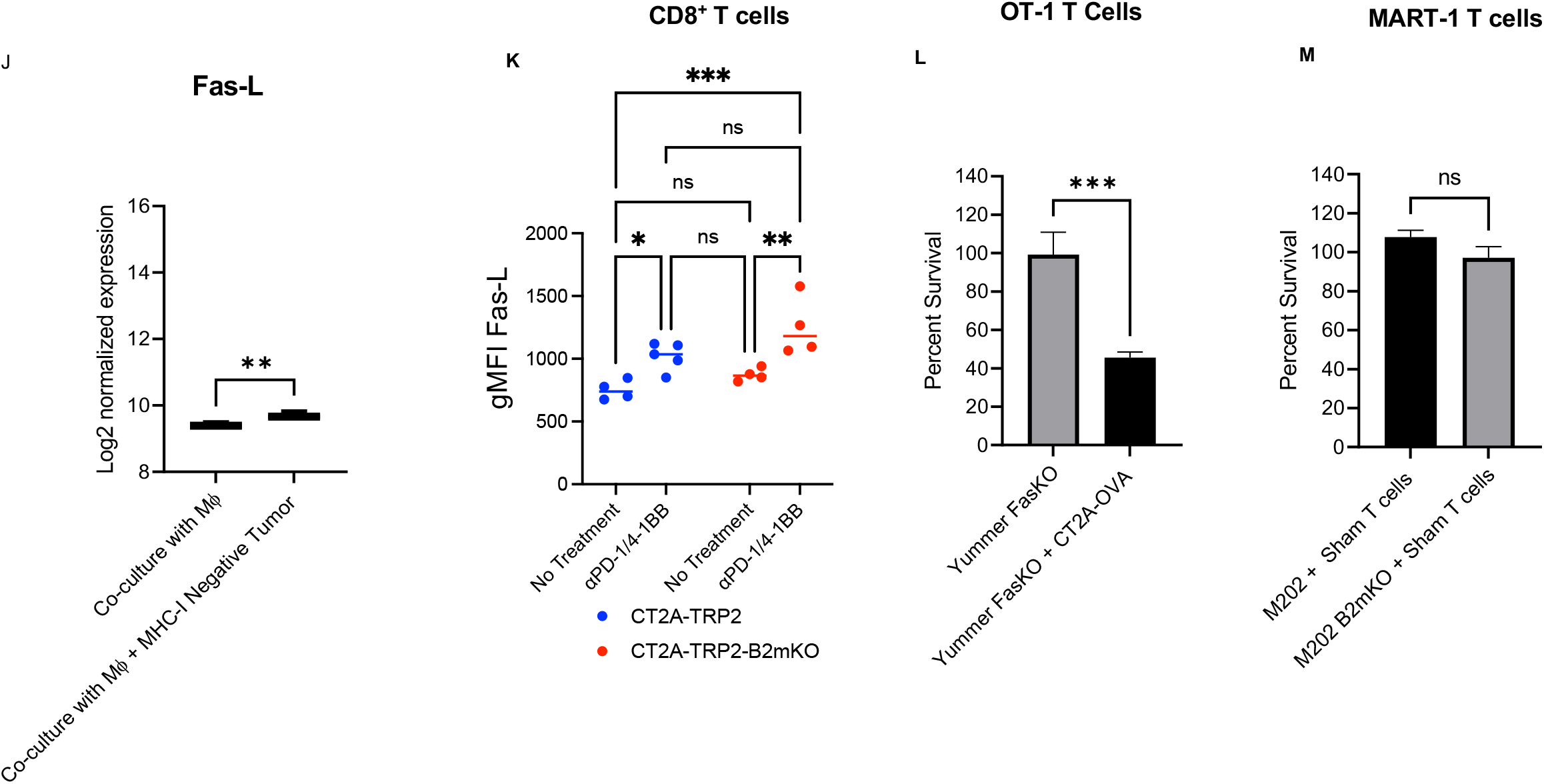
**A-B**. Volcano plot depicting log2 fold change versus log10 adjusted *P* value for differential expression of T cell function genes (**a**) or apoptotic function genes (**b**) between OT-1 T cells co-cultured with OVA macrophages and CT2A-TRP2-β2mKO tumors as compared with OT-1 T cells cultured alone. **C.** Percent survival of CT2A-TRP2-β2mKO cells or CT2A-TRP2 co-cultured with OT-1 CD8^+^ T cells and blocking antibody for TRAIL or isotype control. **D**. Percent NKG2D+ OT-1 CD8^+^ T cells after 12 hours in co-culture with antigen loaded Mϕ or alone in TCM supplemented with IL-2. **E**. CT2A-TRP2-B2mKO expression of MULT-1. **F-g**. GL261-OVA-β2mKO expression of RAE-1 **(f)** or MULT-1 (**g**). **H**. Percent survival of CT2A-OVA in co-culture with OT-1 T cells, CT2A-TRP2-β2mKO tumors and IgG isotype control or αNKG2D. **I-J**. Log2 Normalized expression of granzyme B (**i**) or Fas-L (**j**) in OT-1 T cells co-cultured with cognate antigen loaded macrophages or cognate antigen loaded macrophages and MHC-I-negative CT2A-TRP2-β2mKO. **K**. Geometric mean fluorescence intensity (gMFI) of Fas-L expression in tumor infiltrating CD8^+^ T cells from mice bearing either MHC-I-positive or negative IC tumors +/- treatment with αPD-1/4-1 BB. **L**. Survival of Yummer FasKO cells co-cultured with OT-1 T cells, or OT-1 T cells and OVA+ tumor (OVA stim). **M**. Survival of M202 or M202-β2mKO after 24h co-culture with untransduced human T cells (Sham T cells), 1:1 ET. **O**. Survival of M202 after 24h co-culture with DMF5, M202-β2mKO, and IgG isotype control or αNKG2D, 1:1 ET. *P* values were determined using one-way ANOVA with post-hoc Tukey’s test or by two-tailed, unpaired Student’s t-test.

**Supplementary Table 1.**
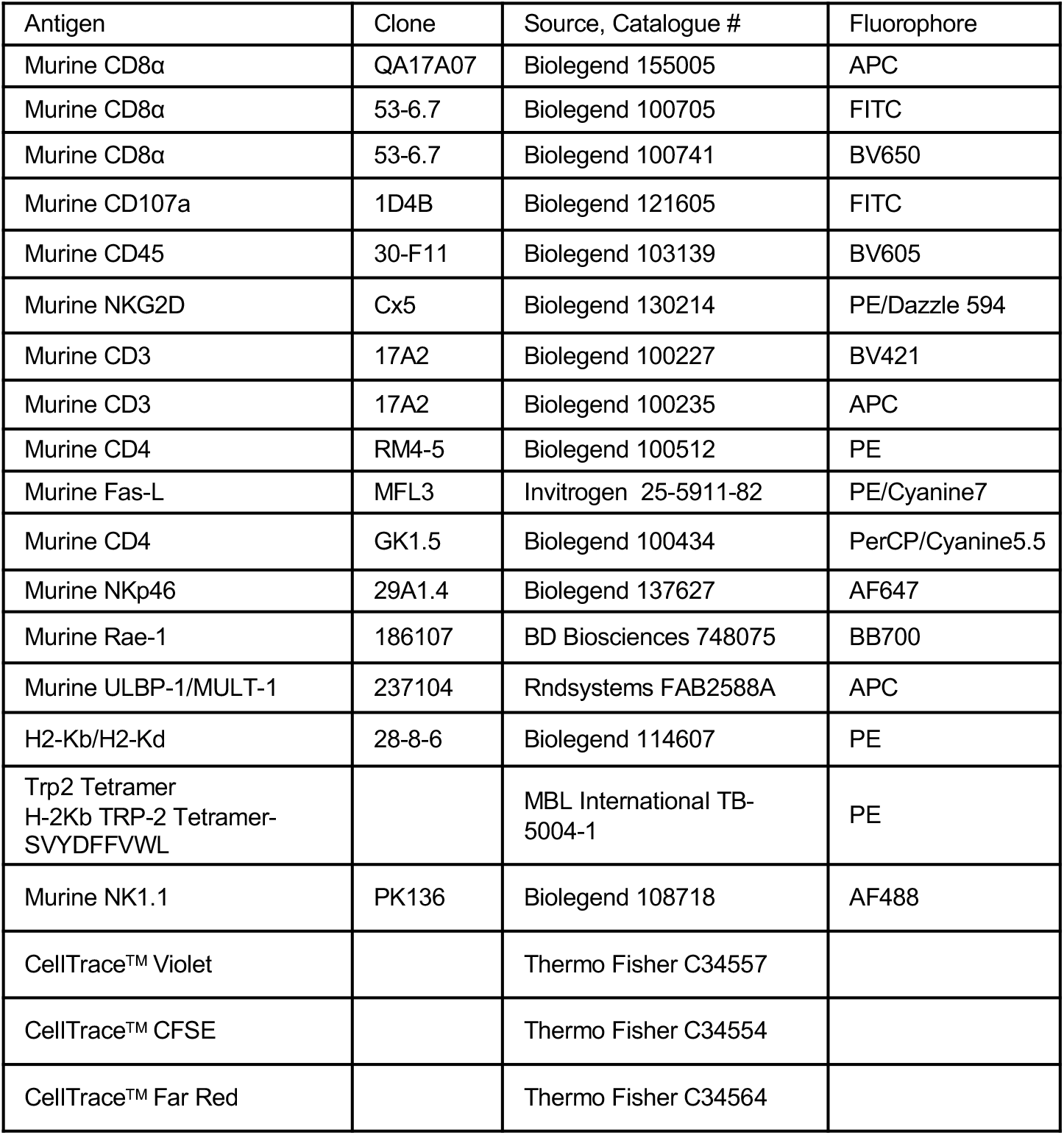
Immunofluorescence antibodies and CellTrace™ Stains used within this manuscript.

## References

1. Cornel, A.M., I.L. Mimpen, and S. Nierkens, MHC Class I Downregulation in Cancer: Underlying Mechanisms and Potential Targets for Cancer Immunotherapy. Cancers (Basel), 2020. 12(7).

2. Zaretsky, J.M., et al., Mutations Associated with Acquired Resistance to PD-1 Blockade in Melanoma. N Engl J Med, 2016. 375(9): p. 819–29.

3. Chisolm, D.A. and A.S. Weinmann, TCR-Signaling Events in Cellular Metabolism and Specialization. Frontiers in Immunology, 2015. 6(292).

4. Bubeník, J., Tumour MHC class I downregulation and immunotherapy (Review). Oncol Rep, 2003. 10(6): p. 2005–2008.

5. Castro M, S.B., Pieper N, Biskup S, Major histocompatibility complex class 1 (MHC1) loss among patients with glioblastoma (GBM). J Clin Oncology, 2020. 38(15).

6. Chow, R.D., et al., AAV-mediated direct in vivo CRISPR screen identifies functional suppressors in glioblastoma. Nature Neuroscience, 2017. 20(10): p. 1329–1341.

7. Gettinger, S., et al., Impaired HLA Class I Antigen Processing and Presentation as a Mechanism of Acquired Resistance to Immune Checkpoint Inhibitors in Lung Cancer. Cancer Discov, 2017. 7(12): p. 1420–1435.

8. Torrejon, D.Y., et al., Overcoming Genetically Based Resistance Mechanisms to PD-1 Blockade. Cancer Discov, 2020. 10(8): p. 1140–1157.

9. Sade-Feldman, M., et al., Resistance to checkpoint blockade therapy through inactivation of antigen presentation. Nat Commun, 2017. 8(1): p. 1136.

10. Busch, E., et al., Beta-2-microglobulin Mutations Are Linked to a Distinct Metastatic Pattern and a Favorable Outcome in Microsatellite-Unstable Stage IV Gastrointestinal Cancers. Front Oncol, 2021. 11: p. 669774.

11. Tang, F., et al., Impact of beta-2 microglobulin expression on the survival of glioma patients via modulating the tumor immune microenvironment. CNS Neurosci Ther, 2021. 27(8): p. 951–962.

12. Woroniecka, K.I., et al., 4-1BB Agonism Averts TIL Exhaustion and Licenses PD-1 Blockade in Glioblastoma and Other Intracranial Cancers. Clin Cancer Res, 2020. 26(6): p. 1349–1358.

13. Woroniecka, K., et al., T Cell Exhaustion Signatures Vary with Tumor Type and are Severe in Glioblastoma. Clin Cancer Res, 2018.

14. Karre, K., et al., Selective rejection of H-2-deficient lymphoma variants suggests alternative immune defence strategy. Nature, 1986. 319(6055): p. 675–8.

15. De Leo, A., A. Ugolini, and F. Veglia, Myeloid Cells in Glioblastoma Microenvironment. Cells, 2020. 10(1).

16. Prajapati, K., et al., Functions of NKG2D in CD8+ T cells: an opportunity for immunotherapy. Cellular & Molecular Immunology, 2018. 15(5): p. 470–479.

17. Markiewicz, M.A., et al., Costimulation through NKG2D Enhances Murine CD8<sup>+</sup> CTL Function: Similarities and Differences between NKG2D and CD28 Costimulation. The Journal of Immunology, 2005. 175(5): p. 2825–2833.

18. Dhar, P. and J.D. Wu, NKG2D and its ligands in cancer. Curr Opin Immunol, 2018. 51: p. 55–61.

19. Raulet, D.H., Roles of the NKG2D immunoreceptor and its ligands. Nature Reviews Immunology, 2003. 3(10): p. 781–790.

20. Venkataraman, G.M., et al., Promoter Region Architecture and Transcriptional Regulation of the Genes for the MHC Class I-Related Chain A and B Ligands of NKG2D. The Journal of Immunology, 2007. 178(2): p. 961–969.

21. Gasser, S. and D.H. Raulet, Activation and self-tolerance of natural killer cells. Immunological Reviews, 2006. 214(1): p. 130–142.

22. Spear, P., et al., NKG2D ligands as therapeutic targets. Cancer Immun, 2013. 13: p. 8.

23. Lanier, L.L., et al., Immunoreceptor DAP12 bearing a tyrosine-based activation motif is involved in activating NK cells. Nature, 1998. 391(6668): p. 703–707.

24. Zompi, S., et al., NKG2D triggers cytotoxicity in mouse NK cells lacking DAP12 or Syk family kinases. Nature Immunology, 2003. 4(6): p. 565–572.

25. Chu, T., et al., Bystander-activated memory CD8 T cells control early pathogen load in an innate-like, NKG2D-dependent manner. Cell Rep, 2013. 3(3): p. 701–8.

26. Billadeau, D.D., et al., NKG2D-DAP10 triggers human NK cell–mediated killing via a Syk-independent regulatory pathway. Nature Immunology, 2003. 4(6): p. 557–564.

27. Groh, V., et al., Fas ligand–mediated paracrine T cell regulation by the receptor NKG2D in tumor immunity. Nature Immunology, 2006. 7(7): p. 755–762.

28. Bauer, S., et al., Activation of NK Cells and T Cells by NKG2D, a Receptor for Stress-Inducible MICA. Science, 1999. 285(5428): p. 727–729.

29. Rincon-Orozco, B., et al., Activation of Vγ9Vδ2 T Cells by NKG2D. The Journal of Immunology, 2005. 175(4): p. 2144–2151.

30. Upadhyay, R., et al., A Critical Role for Fas-Mediated Off-Target Tumor Killing in T-cell Immunotherapy. Cancer Discov, 2021. 11(3): p. 599–613.

31. Upadhyay, R., et al., The death receptor Fas mediates local bystander killing of antigen-negative variants by antigen-specific CD8 T cells in a heterogeneous tumor. The Journal of Immunology, 2019. 202(1 Supplement): p. 138.19–138.19.

32. Smyth, M.J., E. Krasovskis, and R.W. Johnstone, Fas Ligand-Mediated Lysis of Self Bystander Targets by Human Papillomavirus-Specific CD8<sup>+</sup> Cytotoxic T Lymphocytes. Journal of Virology, 1998. 72(7): p. 5948–5954.

33. Johnson, L.A., et al., Gene Transfer of Tumor-Reactive TCR Confers Both High Avidity and Tumor Reactivity to Nonreactive Peripheral Blood Mononuclear Cells and Tumor-Infiltrating Lymphocytes. The Journal of Immunology, 2006. 177(9): p. 6548–6559.

34. Robbins, P.F., et al., Single and Dual Amino Acid Substitutions in TCR CDRs Can Enhance Antigen-Specific T Cell Functions. The Journal of Immunology, 2008. 180(9): p. 6116–6131.

35. Kalbasi, A., et al., Prevention of interleukin-2 withdrawal-induced apoptosis in lymphocytes retrovirally cotransduced with genes encoding an antitumor T-cell receptor and an antiapoptotic protein. Journal of immunotherapy (Hagerstown, Md.: 1997), 2010. 33(7): p. 672–683.

36. Zinkernagel, R.M. and P.C. Doherty, Restriction of in vitro T cell-mediated cytotoxicity in lymphocytic choriomeningitis within a syngeneic or semiallogeneic system. Nature, 1974. 248(5450): p. 701–702.

37. Castro, A., et al., Elevated neoantigen levels in tumors with somatic mutations in the HLA-A, HLA-B, HLA-C and B2M genes. BMC Medical Genomics, 2019. 12(6): p. 107.

38. Sette, A., R. Chesnut, and J. Fikes, HLA expression in cancer: implications for T cell-based immunotherapy. Immunogenetics, 2001. 53(4): p. 255–63.

39. Hicklin, D.J., F.M. Marincola, and S. Ferrone, HLA class I antigen downregulation in human cancers: T-cell immunotherapy revives an old story. Molecular Medicine Today, 1999. 5(4): p. 178–186.

40. Tikidzhieva, A., et al., Microsatellite instability and Beta2-Microglobulin mutations as prognostic markers in colon cancer: results of the FOGT-4 trial. British Journal of Cancer, 2012. 106(6): p. 1239–1245.

41. Koelzer, V.H., et al., Prognostic impact of β-2-microglobulin expression in colorectal cancers stratified by mismatch repair status. Journal of Clinical Pathology, 2012. 65(11): p. 996–1002.

42. Barrow, P., et al., Confirmation that somatic mutations of beta-2 microglobulin correlate with a lack of recurrence in a subset of stage II mismatch repair deficient colorectal cancers from the QUASAR trial. Histopathology, 2019. 75(2): p. 236–246.

43. Ericsson, C., et al., Association of HLA Class I and Class II Antigen Expression and Mortality in Uveal Melanoma. Investigative Ophthalmology & Visual Science, 2001. 42(10): p. 2153–2156.

44. Zhang, H., et al., B2M overexpression correlates with malignancy and immune signatures in human gliomas. Scientific Reports, 2021. 11(1): p. 5045.

45. Middha, S., et al., Majority of B2M-Mutant and-Deficient Colorectal Carcinomas Achieve Clinical Benefit From Immune Checkpoint Inhibitor Therapy and Are Microsatellite Instability-High. JCO Precis Oncol, 2019. 3.

46. Sanjana, N.E., O. Shalem, and F. Zhang, Improved vectors and genome-wide libraries for CRISPR screening. Nat Methods, 2014. 11(8): p. 783–784.

47. Chen, H., et al., NKG2D blockade attenuated cardiac allograft vasculopathy in a mouse model of cardiac transplantation. Clin Exp Immunol, 2013. 173(3): p. 544–52.

48. Bendle, G.M., et al., Lethal graft-versus-host disease in mouse models of T cell receptor gene therapy. Nat Med, 2010. 16(5): p. 565–70, 1p following 570.

49. Nakano, H., et al., Blood-derived inflammatory dendritic cells in lymph nodes stimulate acute T helper type 1 immune responses. Nature immunology, 2009. 10(4): p. 394–402.

50. Huang, M.-N., et al., Antigen-loaded monocyte administration induces potent therapeutic antitumor T cell responses. The Journal of clinical investigation, 2020. 130(2).

51. Tomaszewski, W.H., et al., Broad immunophenotyping of the murine brain tumor microenvironment. Journal of Immunological Methods, 2021. 499: p. 113158.

52. Woroniecka, K., et al., Flow Cytometric Identification of Tumor-Infiltrating Lymphocytes from Glioblastoma. Methods Mol Biol, 2018. 1741: p. 221–226.

53. Chan, K.S. and A. Kaur, Flow cytometric detection of degranulation reveals phenotypic heterogeneity of degranulating CMV-specific CD8+ T lymphocytes in rhesus macaques. J Immunol Methods, 2007. 325(1-2): p. 20–34.

## References

1. Sanjana, N.E., O. Shalem, and F. Zhang, Improved vectors and genome-wide libraries for CRISPR screening. Nat Methods, 2014. 11(8): p. 783–784.

2. Chen, H., et al., NKG2D blockade attenuated cardiac allograft vasculopathy in a mouse model of cardiac transplantation. Clin Exp Immunol, 2013. 173(3): p. 544–52.

3. Bendle, G.M., et al., Lethal graft-versus-host disease in mouse models of T cell receptor gene therapy. Nat Med, 2010. 16(5): p. 565–70, 1p following 570.

4. Johnson, L.A., et al., Gene Transfer of Tumor-Reactive TCR Confers Both High Avidity and Tumor Reactivity to Nonreactive Peripheral Blood Mononuclear Cells and Tumor-Infiltrating Lymphocytes. The Journal of Immunology, 2006. 177(9): p. 6548–6559.

5. Robbins, P.F., et al., Single and Dual Amino Acid Substitutions in TCR CDRs Can Enhance Antigen-Specific T Cell Functions. The Journal of Immunology, 2008. 180(9): p. 6116–6131.

6. Torrejon, D.Y., et al., Overcoming Genetically Based Resistance Mechanisms to PD-1 Blockade. Cancer Discov, 2020. 10(8): p. 1140–1157.

7. Nakano, H., et al., Blood-derived inflammatory dendritic cells in lymph nodes stimulate acute T helper type 1 immune responses. Nature immunology, 2009. 10(4): p. 394–402.

8. Huang, M.-N., et al., Antigen-loaded monocyte administration induces potent therapeutic antitumor T cell responses. The Journal of clinical investigation, 2020. 130(2).

9. Tomaszewski, W.H., et al., Broad immunophenotyping of the murine brain tumor microenvironment. Journal of Immunological Methods, 2021. 499: p. 113158.

10. Woroniecka, K., et al., Flow Cytometric Identification of Tumor-Infiltrating Lymphocytes from Glioblastoma. Methods Mol Biol, 2018. 1741: p. 221–226.

11. Chan, K.S. and A. Kaur, Flow cytometric detection of degranulation reveals phenotypic heterogeneity of degranulating CMV-specific CD8+ T lymphocytes in rhesus macaques. J Immunol Methods, 2007. 325(1-2): p. 20–34.

